# Transcriptional activation of DBP by hnRNP K facilitates circadian rhythm

**DOI:** 10.1101/503771

**Authors:** Paul Kwangho Kwon, Kyung-Ha Lee, Ji-hyung Kim, Sookil Tae, Seokjin Ham, Hyo-Min Kim, Jung-Hyun Choi, Young-Hun Jeong, Sung Wook Kim, Hee Yi, Hyun-Ok Ku, Tae-Young Roh, Chunghun Lim, Kyong-Tai Kim

## Abstract

D-site albumin promoter binding protein (DBP) supports the rhythmic transcription of downstream genes, in part by displaying high-amplitude cycling of its own transcripts compared to other circadian clock genes. However, the underlying mechanism remains elusive. Here, we demonstrated that poly(C) motif within DBP proximal promoters, in addition to an E-box element, provoked the transcriptional activation through increased RNA polymerase 2 (Pol2) recruitment by inducing higher chromatin accessibility. We also clarified that heterogeneous nuclear ribonucleoprotein K (hnRNP K) is a key regulator that binds to the poly(C) motif on single-stranded DNAs in vitro. Chromatin immunoprecipitation further confirmed the expression-dependent and rhythmic binding of hnRNP K which was inhibited through its cytosolic localization mediated by time-dependent ERK activation. Knockdown of hnRNP K triggered low-amplitude mRNA rhythms in DBP and other core clock genes through transcriptional or post-transcriptional regulation. Finally, transgenic depletion of a *Drosophila* homolog of hnRNP K in circadian pacemaker neurons lengthened 24-hour periodicity in free-running locomotor behaviors. Taken together, our results provide new insights into the function of hnRNP K as a transcriptional amplifier of DBP, which acts rhythmically through its intracellular localization by the ERK phosphorylation and as an mRNA stabilizer along with its physiological significance in circadian rhythms of *Drosophila*.

**Significance Statement:** In the case of mood disorders and the aging process, the mRNA expression and amplitude level of clock genes, including DBP, were reported to be diminished. However, the reason behind this decrease of clock gene amplitude and expression level remained unclear. Through this study, we revealed the regulatory mechanism behind the expression of clock genes, especially of DBP mRNA expression. In addition, we discovered that hnRNP K regulates more core clock genes than what we have previously known, such as Clock and Periods. Finally, we demonstrated the physiological significance of hnRNP K in Drosophila through its RNAi line model. Hence, our findings show the regulatory mechanism of circadian rhythm that may provide insight on mood disorder and aging process.

## Introduction

Circadian rhythm is present in living organisms and reflects the elaborate nature of biological systems. Most living organisms have a circadian rhythm, which helps to synchronize daily changes in gene expression, metabolism, and physiology (1). The circadian clock system is modulated by transcriptional, post-transcriptional, translational, and post-translational regulation (2-4). Several studies have reported that clock genes are controlled by core genes such as the BMAL1-CLOCK complex, which acts as an activator (5, 6), and Cryptochrome (Cry1 and Cry2) (7, 8) along with Period (Per1, Per2, and Per3) (9-12) genes, which function as repressors. Recently, post-transcriptional regulation of clock genes by RNA-binding proteins such as hnRNP Q (13-15), PTB (16), hnRNP D (17), hnRNP A1 (18), hnRNP R (19), and hnRNP K (20) was shown to play a role in circadian rhythm.

Disruption of the amplitude of clock gene expression levels has been reported in studies of mood disorders and aging (21-24). Especially, both of D site albumin promoter binding protein (DBP) oscillation amplitude and its expression level were decreased in fibroblasts from bipolar patients (21). Here, DBP mRNA oscillation is known for its high amplitude among clock genes (25, 26). DBP protein has a DNA binding domain that binds specifically to the D-box (TTATG[T/C]AA) and acts as a transcription activator (27-30). In mammalian genomes, there are more than 2,000 putative D-boxes, which govern about 10% of clock-controlled genes (31, 32). The oscillation of DBP mRNA is regulated by the BMAL1-CLOCK complex via multiple E-boxes (4, 33-35). Cry1 also affects the phase of DBP mRNA circadian rhythm (35). However, little is known about the mechanism of high-amplitude DBP mRNA oscillation.

As a strong poly(C)-binding protein (36), heterogeneous nuclear ribonucleoprotein K (hnRNP K) has been revealed to be involved in multiple gene expression processes when bound to single-stranded DNA or RNA, including the regulation of chromatin modification, transcription, post-transcriptional regulation, and translation (37). Among these multiple mechanisms of gene expression regulation, hnRNP K modulates transcription by interacting with TBP (TATA-binding protein) (38, 39) and by stabilizing DNA secondary structures as a trans-activator (40-42).

In this study, we investigated the reason behind high-amplitude DBP mRNA oscillation and the function of hnRNP K to support circadian rhythm in terms of transcriptional regulation. We further validated that the clock function of hnRNP K is conserved in circadian pacemaker neurons of *Drosophila*, where it sustains circadian locomotor behaviors.

## RESULTS

### Oscillation of DBP mRNA expression is supported by its transcription at specific promoter regions

While the high amplitude of DBP mRNA oscillation has been reported, the reason behind the higher amplitude expression of DBP than that of other clock genes remained unclear (25, 26). First, we confirmed the oscillation of clock genes by treating NIH3T3 cells with dexamethasone synchronization for 36 hr and measured the expression level of DBP pre-mRNA and mature mRNA (Figure S1A). In accordance with previous reports, we confirmed that DBP mature mRNA showed a slightly delayed peak time when compared to that of DBP pre-mRNA (43). Next, we measured the oscillation mRNA level of DBP and compared it to that of Rev-erbα and Per2, which are known to be governed by E-box elements. We also compared the oscillation mRNA level of DBP to that of Bmal1, which has an antiphasic rhythm of expression and recognize E-box controlled genes (Figure S1B).

We hypothesized that the major factor that determines the high-amplitude of DBP mRNA expression is transcription efficiency via elevated transcriptional activation. To elucidate potential transcriptional regulatory motifs in eutherian mammals, we aligned the proximal promoter region (genomic regions of 1500 bp from the transcription start site, TSS) of the DBP gene from multiple mammalian species using sequence data in the Ensembl database. We found that short promoter regions (−564 bp from the TSS) were highly alignable among eutherian mammals, suggesting that a possible transcriptional regulatory motif could exist in the proximal promoter region (Figure S1C). We cloned several deletion forms of the proximal promoter region for promoter assays with a luciferase reporter to determine regions critical for transcriptional activation at a peak time of DBP pre-mRNA. Our data indicated that P3 (−400), P4 (−450), P6 (which had lost 209 bp from the TSS, Δ−209), wild-type (−564), and wild-type (Long) (−1500) promoter regions drove significant expression of luciferase, allowing us to determine specific regions of the proximal promoter critical for promoter activity (Figure 1A). In particular, P5 (which we deleted from 209 bp to 450 bp away from TSS, Δ−450/−209) and P2 (−300) constructs had decreased promoter activity relative to that of P3 (−400), P4 (−450), P6 (Δ−209), and wild-type (−564). Next, we identified the oscillation levels of P2, P3, and wild-type using real-time luminescence (Figure 1B). Our data showed that the region between P2 and P3 were necessary for high-amplitude oscillation of DBP transcription. This suggests that other crucial regions for efficient promoter activity exist in the region between 300 bp and 400 bp from the TSS apart from an E-box element. Therefore, we concluded that this specific region of the proximal promoter of DBP could possess critical regions for DBP transcriptional activation.

**Figure 1.**
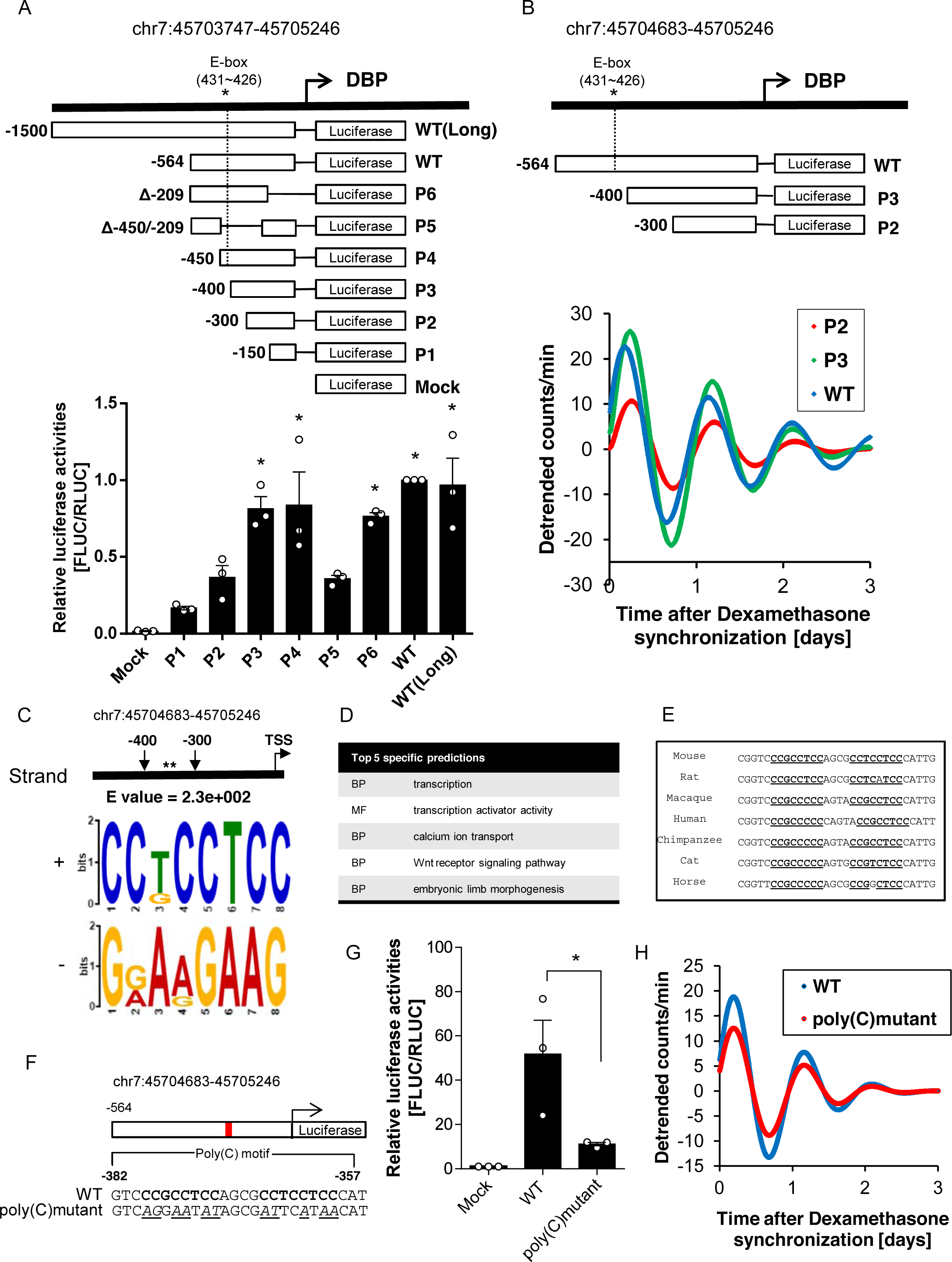
Oscillation of DBP mRNA expression is supported by its transcription at specific promoter regions. (A) Promoter assay was conducted at 24 hrs after Dexamethasone synchronization. P3, P4, P6, WT, and WT(Long) promoter regions showed significant promoter activities, which suggested that the region between −300 and −400 from the TSS was a critical promoter region. In contrast, other promoter regions had relatively little impact on promoter activity. (n=5, indicated by asterisks; * p<0.05 by Tukey’s multiple comparisons test). (B) Representative baseline detrended bioluminescence recordings from WT:LUC, P2:LUC, and P3:LUC, which was transiently transfected into an NIH3T3 cell line that is synchronized by dexamethasone, suggested that the regions between −300 and −400 from the TSS could be crucial in high-amplitude of DBP oscillation. (C) Poly(C) motif (or CCT motif) was scanned in the region between −300 and −400 from TSS using the MEME suite. * indicates the region where poly(C) motif is. (D) One of the predicted functions of poly(C) motif by GOMo was the transcription. Poly(C) motif was the only motif that was predicted to function in transcription among motifs. MF; Molecular function, BP; Biological process. (E) Alignment of genomic regions containing a pair of poly(C) motif showed high conservation of the motif among eutherian mammals, which suggests that a common transcriptional regulatory mechanism is involved in the DBP promoter activity. Underline and bold font represent poly(C) motifs. (F) Illustration of the luciferase reporter system containing WT promoter and poly(C) mutant is described. Bold font indicates poly(C) motif. Underline and italic font represent mutated bases. (G) Promoter activity at 24 hrs after Dexamethasone synchronization of the poly(C) mutant showed a significant decrease compared to WT, which suggests that poly(C) motif is crucial for promoter activity (n=3, Indicated by asterisks; * p<0.05 by Unpaired t-test). (H) Representative baseline detrended bioluminescence recordings of WT:LUC and poly(C) mutant:LUC were measured with luciferase reporter transfected NIH3T3 cell line after dexamethasone synchronization, which showed a decrease in the amplitude level of the poly(C) mutant.

To identify transcriptional regulatory motifs, we used the MEME suite, a tool for searching for specific motifs. We scanned proximal promoter regions with MEME to reveal motifs between P2 (−300) and P3 (−400) (Figure S2A). The poly(C) motif (or CCT motif) (Figure 1C) was the only motif in this region predicted to function in transcription according to the GOMo tool (Figure 1D and Figure S2B).

Furthermore, high-amplitude oscillation of DBP mRNA is often found in mammals such as humans, mice, and rats (21, 27, 44). Therefore, we aligned the poly(C) motif sequences of several eutherian mammals, which revealed that the general transcriptional regulatory mechanism could be mediated by the poly(C) motif (Figure 1E). Next, to identify levels of promoter activity, we cloned a mutated poly(C) motif produced by random substitution from wild-type 5’CCGCCTCCAGCGCCTCCTCC3’ to mutant 5’*AG*G*AA*T*AT*AGCG*AT*TC*A*T*AA*3’ (Figure 1F). We measured the activity of wild-type and poly(C) mutant promoters and found that DBP promoter activity of poly(C) mutant was significantly decreased, which implied that poly(C) motif played a key role in its promoter activity (Figure 1G). Next, we triggered oscillation by synchronizing cells with dexamethasone to identify differences between wild-type and poly(C) mutant promoter activity using real-time luminescence. We found that the amplitude of luciferase expression by the poly(C) mutant was markedly decreased compared to that of wild-type (Figure 1H).

### The poly(C) motif is critical for high-amplitude DBP mRNA oscillation

Next, we designed an NIH3T3 cell line with a deletion of poly(C) motif regions within DBP promoter using the CRISPR-Cas9 system to elucidate the endogenous function of these genomic regions during DBP mRNA oscillation (Figure 2A). After single cell sorting by GFP selection, poly(C) motif regions deleted cells were expanded and confirmed by sequencing in both forward and reverse directions (Figure 2B). We confirmed that the DBP mRNA oscillation amplitude of the poly(C) deleted cells was decreased compared to that of wild-type cells (Figure 2C). We also confirmed that the amplitude of Per 2, which was known to be governed by the D-box element in addition to the E-box element (45), was decreased (Figure 2D). In contrast, Cry1 mRNA oscillation of poly(C) deleted cells were affected slightly when compared to wild-type cells (Figure 2E). Also, Clock mRNA oscillation level was not altered (Figure 2F).

**Figure 2.**
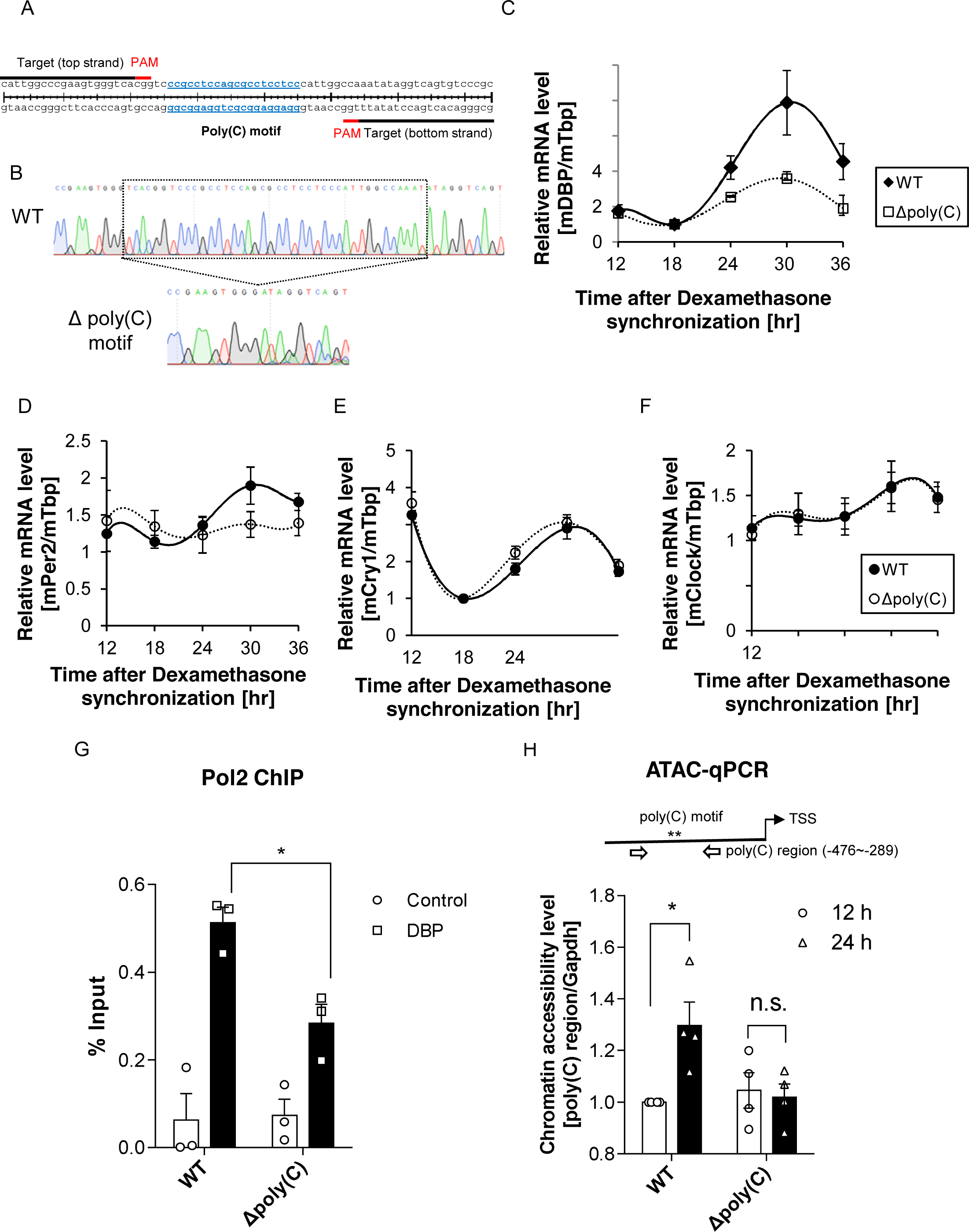
The poly(C) motif is critical for high-amplitude DBP mRNA oscillation. (A) Schematic description of a pair of sgRNAs designed to delete endogenous poly(C) motif regions within the DBP proximal promoter region is shown. (B) Poly(C) motif region deleted cell line made through CRISPR-Cas9 system was confirmed by sequencing. (C) The oscillation of endogenous DBP mRNA was quantified and the poly(C) motif region deleted cell line showed a decrease in amplitude of DBP mRNA oscillation (n=4, Error bar: S.E.M.). (D) Per2 mRNA oscillation level was quantified, which oscillation was affected by the poly(C) motif within the DBP promoter (n=5, Error bar: S.E.M.). (E) Cry1 mRNA oscillation was measured. The phase of Cry1 mRNA oscillation was slightly changed (n=4, Error bar: S.E.M.). (F) Clock mRNA oscillation was used as a control (n=5, Error bar: S.E.M.). (G) Pol2 ChIP was conducted to measure the recruitment level of Pol2 to DBP promoter regions at the peak time of DBP mRNA level (n=3, indicated by asterisks; * p<0.05 by Tukey’s multiple comparisons test). (H) ATAC-qPCR data indicated that chromatin accessibility of WT cells increased at 24 hrs after Dexamethasone synchronization, however, poly(C) motif region deleted cells did not show an increase in the chromatin accessibility (n=4, n.s.: not significant, indicated by asterisks; * p<0.05 by Tukey’s multiple comparisons test).

To identify transcriptional regulation on DBP at the peak time, we measured the level of RNA Polymerase 2 (Pol2) binding during the peak time of DBP pre-mRNA oscillation. Control region of Pol2 ChIP was previously reported (46). The data showed that Pol2 binding level on the DBP promoter region was decreased in poly(C) deleted cells (Figure 2G).

Finally, to clarify the reason behind the reduced pol2 binding level on DBP promoter of poly(C) deleted cells, we utilized the Assay for Transposase-Accessible Chromatin (ATAC) along with real-time PCR. Interestingly, ATAC-qPCR data showed that chromatin accessibility levels were regulated by poly(C) motif region depending on the circadian rhythm (Figure 2H). These results propose that DBP mRNA oscillation might be facilitated by the modulation of chromatin accessibility mediated by poly(C) motif regions.

### hnRNP K binds to the single-stranded poly(C) motif directly and rhythmically

To demonstrate the poly(C) motif binding factor, we pulled down proteins that were bound onto poly(C) oligonucleotide. Descriptions of the forward and reverse poly(C) motif sequences are provided in Figure 3A. We attached biotin to the 5’ nucleotide (Solgent, South Korea) of the forward sequence 5’CCGCCTCCAGCGCCTCCTCC3’ and the reverse complementary sequence 5’GGAGGAGGCGCTGGAGGCGG3’ and performed pull-down experiments with streptavidin in HEK293A cells. We found that almost no proteins were pulled down with the duplex form of the oligonucleotide (data not shown). Thus, we presumed that the single strand oligonucleotide of the poly(C) motif might interact with certain unknown transcription factors. Surprisingly, we identified that a single, thick protein band was detected when we used the forward poly(C) motif single-strand oligonucleotide, which was identified as hnRNP K by mass spectrometry (Figure 3A, Figure S3A). The CCT element is known to be a target of hnRNP K, a single-stranded DNA binding protein that functions as a regulator of the expression of multiple genes (37, 38, 40, 41). We also detected the binding of hnRNP K using anti-hnRNP K at the poly(C) motif (Figure S3B).

**Figure 3.**
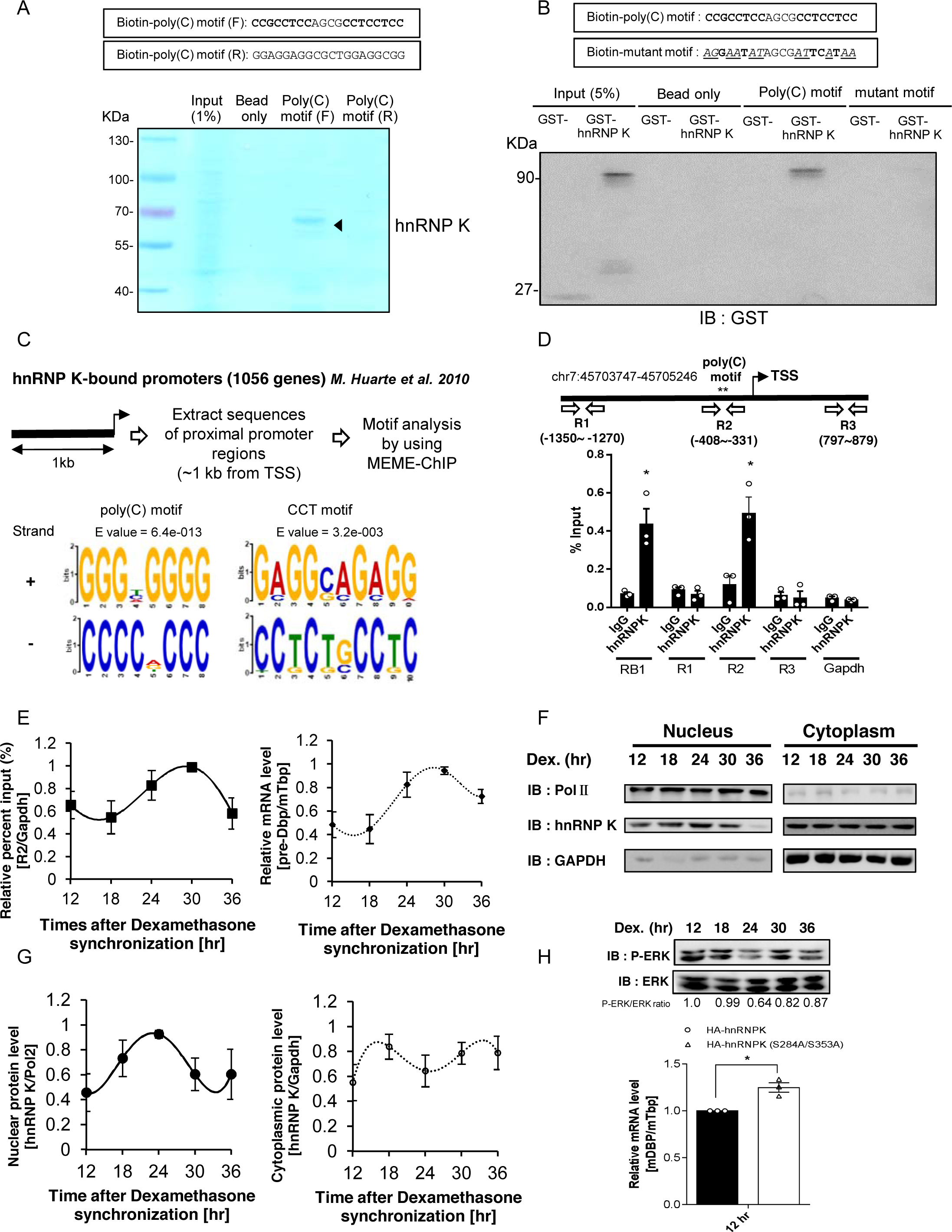
hnRNP K binds to single-stranded poly(C) motif directly and rhythmically. (A) Coomassie staining showed that one thick band was bound onto the forward poly(C) motif oligonucleotide, which was identified as hnRNP K by Mass spectrometry. In contrast, there were no prominent proteins detected at complementary poly(C) motif oligonucleotide. (B) Purification of GST and GST-hnRNP K was also carried out for GST pull-down assay. GST-hnRNP K was detected at wild-type oligonucleotide, but not at the mutant oligonucleotide, which demonstrates that hnRNP K is directly bound to poly(C) motif. (C) The schematic description of the motif analysis is shown. Proximal promoter regions of 1056 genes were analyzed using the database from *M. Huarte et al. 2010.* Motif analysis showed their poly(C) motif was frequently found along with a CCT motif. (D) Endogenous binding of hnRNP K on proximal promoter regions of DBP was confirmed by using ChIP assay. R1, R2, and R3 indicate each genomic regions. RB1 was used as a positive control, while Gapdh was used as a negative control (n=3, indicated by asterisks; * p<0.05 by Tukey’s multiple comparisons test). (E) Rhythmic binding of hnRNP K was confirmed through triggering mRNA oscillation in NIH3T3 cells. Pre-mRNA levels of DBP showed the proportional relationship between transcription of DBP and the binding levels of hnRNP K. Relative percent input (%) indicates the percent input (%) level normalized by the Gapdh percent input (%) level (n=3, Error bar: S.E.M.). (F) The rhythmic expression of the nuclear hnRNP K protein could be a major contributing factor of rhythmic binding of hnRNP K. After triggering the circadian rhythm by dexamethasone synchronization, the levels of nuclear hnRNP K and cytoplasmic hnRNP K were measured after the subcellular fractionation. Pol2 was used as a nuclear marker gene and GAPDH was used as a cytoplasmic marker gene. (G) The nuclear protein levels and cytoplasmic protein levels of hnRNP K were quantified (n=5, Error bar: S.E.M.). (H) Wild-type and phosphomutant hnRNP K overexpression before times after Dexamethasone synchronization indicated that mutant hnRNP K raised DBP mRNA more than wild-type hnRNP K at the peak time of ERK activation level (n=3, Indicated by asterisks; * p<0.05 by Unpaired t-test).

To confirm direct binding of hnRNP K to the poly(C) motif oligonucleotide, we purified GST and GST-hnRNP K proteins and performed pull-down experiments with 5’ biotinylated poly(C) motif and 5’ biotinylated poly(C) mutant (Figure 3B). We confirmed that GST-hnRNP K was detected only in the wild-type oligonucleotide pulled-down group through immunoblotting (Figure 3B). This demonstrated that hnRNP K binds directly to the poly(C) motif of DBP promoter regions *in vitro.*

Although the CCT element has already been reported to be a binding target of hnRNP K, functional confirmation of this interaction and prediction of the function of hnRNP K on this motif has not previously been investigated (37, 38, 40, 41). For this reason, we searched a database that contains 1056 hnRNP K–bound promoter genes from hnRNP K ChIP-Seq (47) for specific motifs that were predicted to function in transcription using MEME-ChIP (Figure S3C). We detected poly(C) and CCT motifs within the proximal promoter regions of hnRNP K–bound promoter genes (Figure 3C), both of which are predicted to function as transcription (Figure S3C). This suggests that hnRNP K binds to single-stranded poly(C) and CCT motifs and may, therefore, function as a transcription activator.

Next, we analyzed the KH3 domain sequence of hnRNP K (Figure S4A), which is a single-stranded DNA binding domain (48-50). Using the Ensembl database, we found that the KH3 domain of several eutherian mammals aligned perfectly (Figure S4B). This result suggests that hnRNP K binding to the promoter region of DBP could be a critical regulatory mechanism for DBP transcription in eutherian mammals.

Previously, we confirmed that hnRNP K bound to the poly(C) motif *in vitro* and predicted that this would allow hnRNP K to function as a transcriptional activator using Gene Ontology for Motifs (GOMo). We then examined the endogenous binding of hnRNP K to the proximal promoter region of DBP using a Chromatin Immunoprecipitation (ChIP) assay, because previously reported ChIP-seq data did not show DBP as a target gene of hnRNP K. The ChIP results showed that hnRNP K specifically bound to Region 2 (R2), which contains a poly(C) motif between −300 and −400 from the TSS. Region 1 (R1) and Region 3 (R3) were not bound by hnRNP K (Figure 3D). The RB1 gene, to which hnRNP K binds robustly, was used as a positive control, while Gapdh was used as a negative control (47). Additionally, we investigated whether the binding of hnRNP K was rhythmically influenced by dexamethasone synchronization, which induces mRNA circadian oscillation of clock genes, to further clarify the involvement of hnRNP K in DBP transcription. We found that oscillation of DBP pre-mRNA levels correlated with the rhythmic binding of hnRNP K to R2 (Figure 3E). Furthermore, we confirmed the subcellular protein expression levels of hnRNP K by subcellular fractionation of NIH3T3 cells. These data suggest that the rhythmic expression of nuclear hnRNP K proteins may be a major contributing factor to the rhythmic binding of hnRNP K (Figure 3F and 3G).

Because it was previously reported that hnRNP K transport was controlled by an ERK activation and that ERK activation was mediated in rhythmic manners, we demonstrated the time-dependent transport of hnRNP K along with activation level of ERK (51-53). Interestingly, we identified that the peak time of ERK activation level was inversely related with the peak time of hnRNP K nuclear expression level after dexamethasone synchronization (Figure 3H). Next, we overexpressed hnRNP K WT and hnRNP K phosphomutant (which was a direct target site of phosphorylation by ERK activation) at both the peak time (Figure 3H) and trough time (Figure S4C) of ERK phosphorylation. The result showed that phosphomutant hnRNP K raised DBP mRNA levels at the peak time of ERK activation, but not the wildtype hnRNP K, which meant that phosphorylation of hnRNP K by ERK activation inhibited transcription in the nucleus by transporting hnRNP K to the cytoplasm.

Thus, our data suggest that time-dependent ERK activation controls hnRNP K transport, which regulates the binding of hnRNP K onto poly(C) motif regions resulting in the robust oscillation of DBP mRNA. This finding led us to explore possible mechanisms of high-amplitude DBP mRNA expression.

### hnRNP K expression level controls the transcription of DBP through the poly(C) motif

We then assessed promoter activity through knockdown of hnRNP K. The knockdown efficiency of the hnRNP K siRNA pool was measured by immunoblotting (Figure 4A). Cells transfected with the hnRNP K siRNA pool showed an approximately 70% decrease in hnRNP K protein levels than the siRNA-transfected control group. Furthermore, we checked cell viability using the MTT assay. Under our experimental conditions, knockdown of hnRNP K did not significantly affect cell viability (Figure 4B).

**Figure 4.**
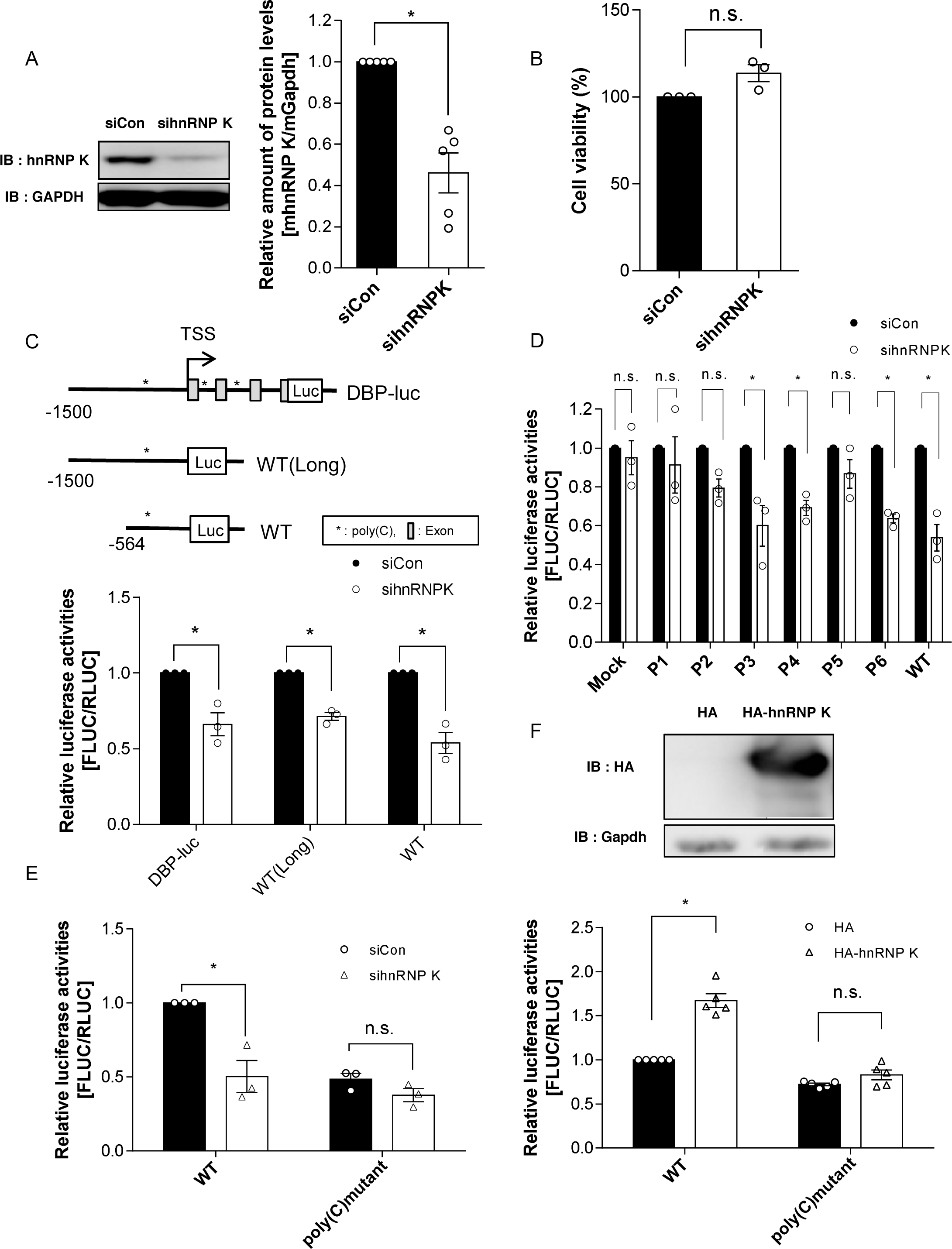
hnRNP K expression level controls the transcription of DBP through the poly(C) motif. (A) Knockdown of hnRNP K was mediated by siRNA transfection in NIH3T3 cells. The group with hnRNP K siRNA pool transfection showed a significant decrease of the hnRNP K protein level (n=5, indicated by asterisks; * p<0.05 by Unpaired t-test). (B) MTT assay showed that the knockdown of hnRNP K did not significantly affect the cell viability of NIH3T3 cell line (n=3, n.s., not significant; p<0.05 by Unpaired t-test). (C) Along with WT and WT(Long), the level of DBP-luciferase which contained regulatory motifs within exons and introns was similarly downregulated by the lack of hnRNP K (n=3, indicated by asterisks; * p<0.05 by Sidak’s multiple comparisons test). (D) Knockdown of hnRNP K decreased the level of P3, P4, P6, and WT promoter activity which contained the poly(C) motif regions (n=3, n.s.: not significant, indicated by asterisks; * p<0.05 by Tukey’s multiple comparisons test). (E) To identify the effect of knockdown of hnRNP K on WT and poly(C) mutant, promoter assay was conducted with siRNA transfection. It showed a significant decrease only with WT promoter activity (n=3, n.s.: not significant, indicated by asterisks; * p<0.05 by One-way ANOVA). (F) Overexpression of hnRNP K increased a WT promoter activity, whereas the promoter activity of the mutant was not altered (n=5, n.s.: not significant, indicated by asterisks; * p<0.05 by One-way ANOVA).

We compared the levels of DBP-luciferase constructs containing all regulatory elements including the poly(C) motif, along with wild-type and wild-type (Long) constructs, during knockdown of hnRNP K. We confirmed that levels of DBP-luciferase, wild-type, and wild-type (Long) expression were down-regulated by knockdown of hnRNP K (Figure 4C). Next, we measured the promoter activity of constructs that contained serially deleted proximal promoter regions as shown in Figure 1A. We also confirmed that the promoter activities of the P3, P4, P6, and wild-type constructs decreased significantly when hnRNP K was downregulated (Figure 4D). A poly(C) motif was present in the P3, P4, P6, and wild-type promoters, which implies a correlation between the poly(C) motif and hnRNP K. Next, we conducted a promoter assay with the wild-type and poly(C) mutant shown in Figure 1F during knockdown of hnRNP K. While this clearly decreased the promoter activity of the wild-type, the promoter activity of the poly(C) mutant changed only a little (Figure 4E). We also overexpressed hnRNP K using a human influenza hemagglutinin (HA) tag to assess promoter activity. Overexpression of hnRNP K increased wild-type promoter activity only (Figure 4F). This indicates that hnRNP K could function as a transcriptional activator by binding to the poly(C) motif in the DBP promoter.

### hnRNP K supports high-amplitude DBP oscillation by influencing transcription, not mRNA stability

After clarifying DBP promoter activity through luciferase reporter assays, we measured the endogenous levels of DBP pre-mRNA and mature mRNA. Levels of both these types of DBP mRNA decreased after knockdown of hnRNP K (Figure 5A). Per3, which has been reported to retain mRNA stability through hnRNP K, was used as a positive control (20). In addition, we elucidated the kinetics of mRNA decay to exclude the possibility of mRNA degradation, which was previously reported to be related to the function of hnRNP K (20). To examine the role of hnRNP K in DBP regulation, the mRNA stability of the control group and the siRNA knockdown group were compared under treatment with actinomycin D, which inhibits transcription. The half-life of DBP mRNA did not change during the knockdown of hnRNP K (Figure 5B). This indicates that hnRNP K regulates DBP not by mRNA degradation, but by transcriptional activation.

**Figure 5.**
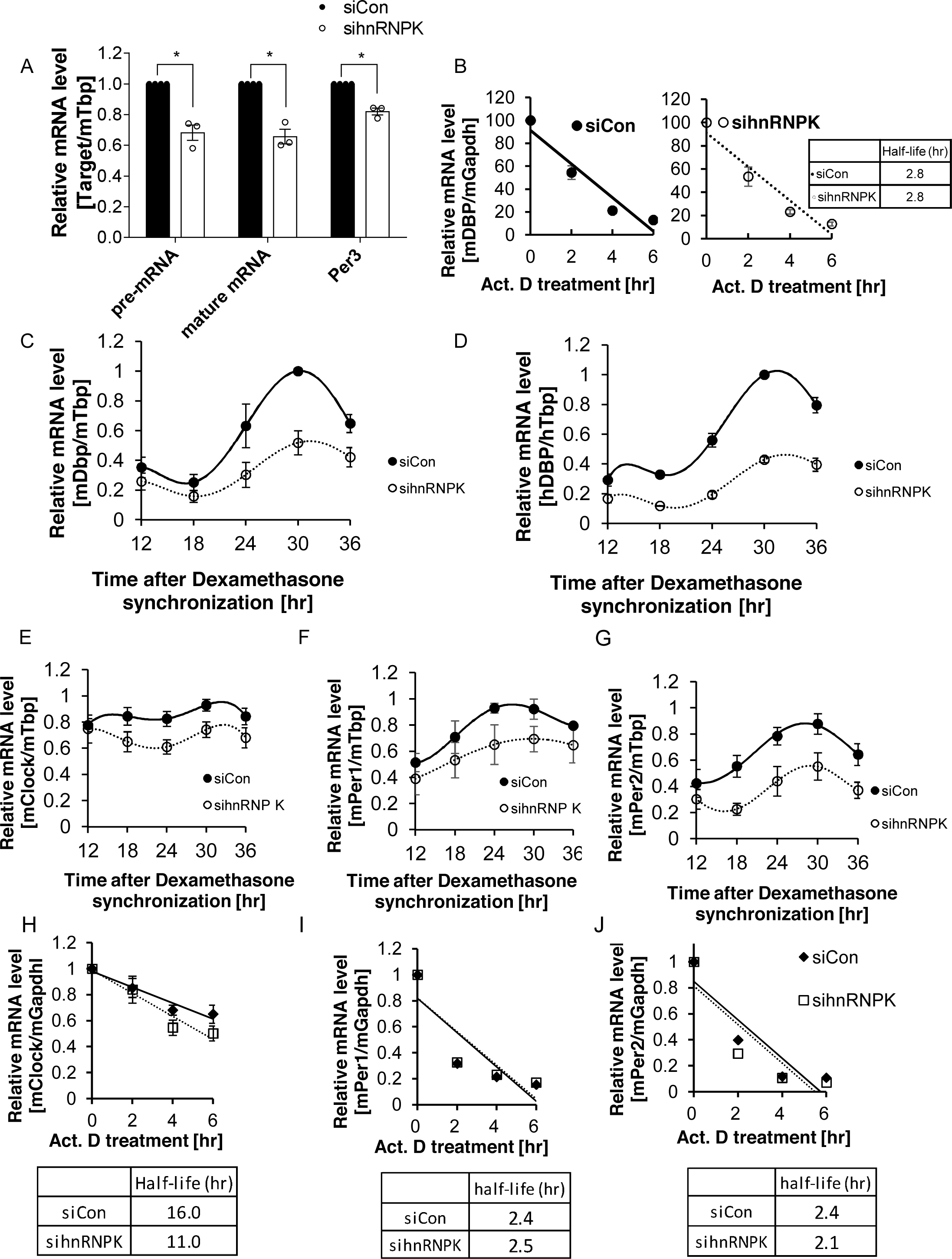
hnRNP K supports high-amplitude DBP oscillation by influencing transcription, not mRNA stability. (A) Pre-mRNA and mature mRNA levels of DBP were measured during knockdown of hnRNP K. Both mRNA levels of DBP were decreased. Per3 was used as a positive control. (n=3, indicated by asterisks; * p<0.05 by Unpaired t-test). (B) During knockdown of hnRNP K, mRNA decay rates of DBP was not altered when transcription inhibitor, actinomycin D, was treated (‘siCon’ group: y = −14.71x + 91.323, ‘sihnRNP K’ group: y = −14.603x + 91.192, n=3, Error bar: S.E.M.). (C) Oscil-lation of DBP mRNA of mouse NIH3T3 cells was measured after dexamethasone synchronization. hnRNP K knockdown group showed a decrease in mRNA amplitude of DBP (n=3, Error bar: S.E.M.). (D) Dexamethasone synchronized human U2OS cells also showed a decrease in amplitude of DBP during the knockdown of hnRNP K (n=3, Error bar: S.E.M.). (E-G) Per1, Per2, and, Clock mRNA oscillation levels were quantified (n=3, Error bar: S.E.M.). (H-J) Clock (‘siCon’ group: y = −6.0883x + 97.896, ‘sihnRNP K’ group: y = −8.9137x + 98.99, n=3, Error bar: S.E.M.), Per1 (‘siCon’ group: y = - 0.1319x + 0.8176, ‘sihnRNP K’ group: y = −0.1291x + 0.8196, n=4, Error bar: S.E.M.), and Per2 (‘siCon’ group: y = −0.1478x + 0.8497, ‘sihnRNP K’ group: y = −0.1486x + 0.8144, n=4, Error bar: S.E.M.) mRNA decay kinetics were measured. It indicated that Clock mRNA stability might be controlled by hnRNP K, however, Per1 and Per2 mRNA were not or marginal.

The circadian rhythm of DBP mRNA in mouse NIH3T3 cells was also measured after triggering oscillation with dexamethasone to clarify the role of hnRNP K in maintaining high-amplitude DBP mRNA oscillation. Quantification of circadian rhythm oscillation under hnRNP K knockdown revealed a decrease in DBP mRNA oscillation amplitude (Figure 5C). This result indicates that high-amplitude DBP oscillations are maintained through transcriptional activation by hnRNP K.

Next, we examined the effects of hnRNP K on other D-box trans-acting regulators—thyrotroph embryonic factor (Tef), hepatic leukemia factor (Hlf), and nuclear factor interleukin 3 regulated (Nfil3)—by measuring mRNA levels during knockdown of hnRNP K. We confirmed that the mRNA level of DBP decreased significantly during knockdown of hnRNP K, while mRNA levels of Tef, Hlf, and Nfil3 did not (Figure S5A). This indicates that the primary target of hnRNP K is the D-box regulator DBP, not other D-box regulators.

Previously, we revealed that poly(C) motif regions among eutherian mammals could function as the binding site of transcriptional activators. We found that the mRNA level of hnRNP K in human U2OS cells was 30% of that of the control when hnRNP K was knocked down (Figure S5B). Furthermore, cell viability testing by the MTT assay demonstrated that knockdown of hnRNP K did not significantly affect cell viability under our experimental conditions (Figure S5C). Oscillation amplitude of human DBP mRNA was also decreased during hnRNP K knockdown (Figure 5D).

Finally, we investigated the mRNA levels of other core clock genes, including Bmal1, Clock, Cry1, Cry2, Period1, and Period2 during the knockdown of hnRNP K. Interestingly, even though the mRNA level of Bmal1, Cry1, and Cry2 were not changed significantly, mRNA levels of Clock, Period1 and Period 2 were down-regulated during knockdown of hnRNP K (Figure S5D). Next, we confirmed that mRNA oscillation levels of Per1, Per2, and, Clock. We found that mRNA oscillation levels of Per1, Per2, and Clock were lowered after Dexamethasone synchronization during knockdown of hnRNP K (Figure 5E-5G). In contrast, Cry1 mRNA oscillation level was only slightly changed (Figure S5E). Also, Bmal1 mRNA was only marginally lowered during the knockdown of hnRNP K (Figure S5F). Interestingly, we measured mRNA decay kinetics of other clock genes, which demonstrated that Clock might be controlled post-transcriptionally by hnRNP K, while Per1 and Per2 showed marginal difference (Figure 5H-5J).

In conclusion, we propose that DBP mRNA oscillation, which is mediated by hnRNP K, also affects other core clock genes such as Clock and Periods, transcriptionally and post-transcriptionally. This leads us to assume that the rhythmic behaviors of an organism could be affected during the knockdown of hnRNP K.

### hnRNP K depletion in *Drosophila* circadian pacemaker neurons impairs circadian behaviors

hnRNP K homologs are well conserved in other species including *Drosophila* (37). Moreover, it has been demonstrated that *Drosophila* hnRNP K proteins associate selectively with poly(C) nucleotides (54, 55). We, therefore, took a genetic approach in *Drosophila* to determine if hnRNP K-dependent clock function is conserved in non-mammalian species and sustains circadian rhythm at the organismal level. To this end, we quantitatively analyzed circadian locomotor activities in individual flies as a readout of the robustness of circadian rhythms.

Since a strong hypomorphic mutation in the *Drosophila* hnRNP K gene causes lethality (55, 56), we expressed an RNA interference transgene against hnRNP K in different groups of circadian pacemaker neurons and examined its effects on circadian locomotor rhythms in adult flies. We first found that hnRNP K depletion in a small number of circadian pacemaker neurons called posterior dorsal neurons 1 (DN1p) (57-60) (by R18H11-Gal4) suppressed evening activity peak anticipatory to lights-off in 12-hour light: 12-hour dark (LD) cycles (Figure 6A, top). Transgenic flies were further transferred from LD cycles to constant dark (DD) where their rhythmic locomotor activities were primarily driven by endogenous circadian clocks. In the first DD cycle, the hnRNP K depletion evidently delayed the circadian phases of morning and evening activity peaks (Figure 6A, middle), leading to long-period rhythms in the subsequent DD cycles (Figure 6A, bottom). Behavioral analyses using three independent RNAi transgenes consistently validated that hnRNP K depletion in DN1p neurons lengthened the circadian periodicity in DD locomotor rhythms by ~2 hours (Figure 6B) while weaker rhythmicity was observed only in one of the three RNAi lines (Figure 6C).

**Figure 6.**
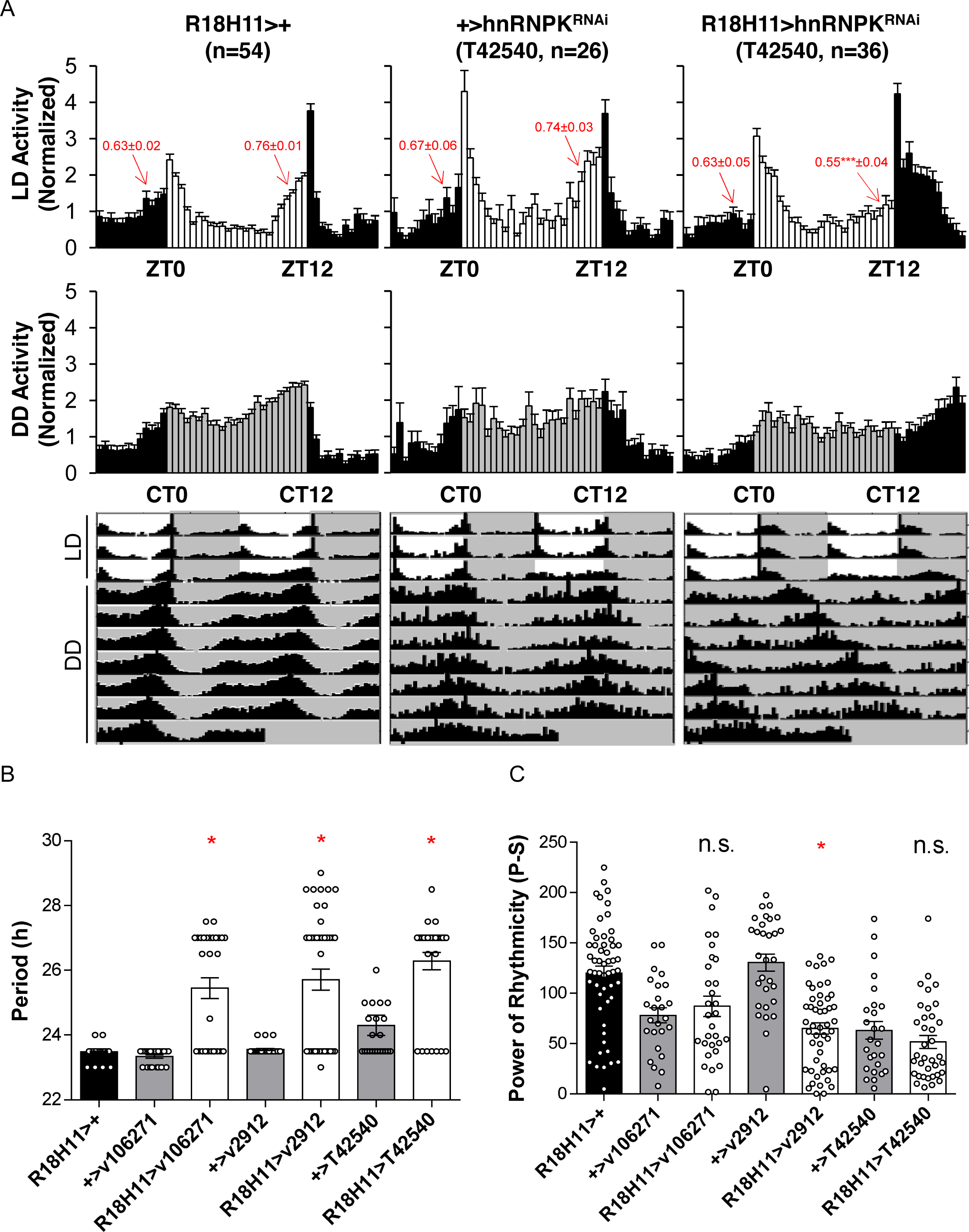
*Drosophila* hnRNP K in DN1p neurons sustains circadian behaviors. (A) hnRNP K depletion in DN1p neurons affected circadian locomotor activities in both LD (12-hour light: 12-hour dark) and DD (constant dark) cycles. Normalized activity profiles in LD cycles (top) or in the first DD cycle (middle) were averaged from individual male flies (n=26-54) while their averaged actograms were double-plotted (bottom). Arrows indicate gradual increases in locomotor activities anticipatory to lights-on or lights-off, which were quantified by morning or evening index values, respectively (average +/- SEM). ZT, zeitgeber time; CT, circadian time; white/black bars, LD cycles; gray/black bars, DD cycles. (Error bar: S.E.M.). (B) Circadian periods in DD locomotor rhythms were averaged from rhythmic flies (*P*-*S*>10) (n=24-53). (C) The power of rhythmicity in DD locomotor rhythms was determined by the chisquared periodograms of individual flies and were averaged per each genotype (n=26-54) (n.s., not significant; \**P* < 0.05 to both heterozygous controls as determined by one-way ANOVA, Tukey post hoc test. Error bar: S.E.M.).

By contrast, we could not observe any comparable phenotypes in circadian behaviors when hnRNP K was depleted in either circadian pacemaker cells broadly expressing TIMELESS (by *tim*-Gal4) (61) or lateral neurons expressing a circadian neuropeptide PIGMENT-DISPERSING FACTOR (by *Pdf*-Gal4) (62) (Figure S6A and S6B). We reason that possible differences in relative abundance of hnRNP K expression and/or efficiency of its transgenic depletion among different groups of circadian pacemaker neurons might explain the long-period rhythms caused selectively by the strong DN1p Gal4 driver. Alternatively, but not exclusively, DN1p neurons might specifically express a clock-relevant gene of which the expression is regulated by hnRNP K and thereby limiting the 24-hour periodicity in DD locomotor rhythms of hnRNP K-depleted flies. Nonetheless, these data demonstrate that hnRNP K homologs sustain circadian rhythms in both flies and mammals, although genetic or neural substrates of hnRNP K-dependent clocks might be diverse.

## DISCUSSION

One of the main findings in this study is that high-amplitude DBP mRNA oscillations are mediated by hnRNP K, which functions as a transcriptional activator. Prior to this study, it was reported that the Bmal1-Clock complex is the main regulator of rhythmic DBP mRNA expression via multiple E-boxes (4, 33-35). Here, we unveiled an additional and novel regulating mechanism that maintains high-amplitude oscillation. A recent study measured rhythmic H3K27ac enrichment and Pol2 density within the DBP proximal promoter regions using ChIP-seq (63). Together with our study findings, this suggests that DBP rhythmic transcription is regulated through a combined regulatory mechanism involving hnRNP K. We demonstrated that hnRNP K could indeed affect the circadian rhythm oscillation of DBP mRNA by acting as a transcription activator along with other complex regulating mechanisms.

Although the importance of maintaining the amplitude of clock gene oscillations has been established, the mechanisms underlying this have remained unclear. Therefore, we investigated the oscillation pattern of DBP and compared it to that of other clock genes (Figure S1A and S1B). Next, we aligned the genomic region upstream of the TSS of DBP among eutherian mammals to determine transcriptional regulatory motifs (Figure S1C). Analysis of the promoter alignment using the MEME suite revealed a poly(C) motif that was predicted to be a transcriptional activator. We confirmed the promoter activity and function of the poly(C) motif using luciferase reporter assays and the CRISPR-Cas9 system (Figure 2). At the same time, we confirmed the presence of the poly(C) motif in proximal promoter regions by analyzing hnRNP K ChIP-ChIP data using MEME-ChIP (47, 64) (Figure 3C). This novel approach towards motif analysis provides a systematic way to confirm predictions experimentally.

hnRNP K binds to single-stranded DNA to stabilize the secondary structure of DNA, thereby promoting transcription (42). Here, we determined hnRNP K, known to be present in yeast to mammals (36, 37), to be the binding partner of the poly(C) motif using an oligonucleotide pulldown assay (Figure 3A and 3B). We also confirmed the binding of endogenous hnRNP K to the poly(C) motif in the DBP proximal promoter region, as well as its rhythmic interaction during circadian oscillation (Figure 3D and 3E). This suggests that the poly(C) motif and hnRNP K might function as critical regulators of circadian rhythm in eutherian mammals. Moreover, we found that the regulation of DBP promoter activity was dependent on the level of nuclear hnRNP K (Figure 4). Also, hnRNP K transport level by ERK activation, in addition to its protein expression levels, could affect its rhythmic binding to the poly(C) motif, resulting in high-amplitude DBP oscillation (Figure 3H).

Furthermore, we measured the endogenous DBP mRNA level after knockdown of hnRNP K (Figure 5A). We demonstrated a decrease in the amplitude of DBP mRNA oscillation by suppressing the level of hnRNP K in mouse and human cells (Figure 5C and 5D). We, therefore, suggest that circadian rhythm is maintained by the level of hnRNP K. In summary, we demonstrated hnRNP K-mediated transcriptional regulation of DBP, which is critical for the robust maintenance of high amplitude DBP mRNA oscillations.

This study provides the first evidence that the relationship between hnRNP K and DBP is crucial for transcription and circadian rhythm. In addition to this, we verified mRNA oscillations of other core clock genes that are controlled by post-transcriptional or transcriptional regulation during low-level of hnRNP K (Figure 5H-5J). Here, we investigated the mechanistic relationship between hnRNP K and DBP that results in delicate control of circadian rhythm in complicated biological systems. Although the precise mechanisms of rhythmic interaction and the involvement of hnRNP K in rhythmic expression remain to be identified, we showed that hnRNP K–mediated transcription and post-transcriptional regulation impacts the oscillation of clock genes and proposed conservation of hnRNP K-dependent clocks.

## MATERIAL AND METHODS

### Cell culture and drug treatment

NIH 3T3 cells, HEK293A, and U2OS cells were cultured in Dulbecco’s modified Eagle’s medium (HyClone) supplemented with 10% fetal bovine serum (HyClone) and 1% antibiotics (WelGENE) and were maintained in a humidified incubator with 95% air and 5% CO2. NIH 3T3 and U2OS cells were synchronized by treatment with 100nM dexamethasone in 12-well plates containing ~3.0×10^5^ cells per well. After 2 hrs, the medium was replaced with complete medium and the cells were harvested every 6 hrs for each sample (15). To measure mRNA decay kinetics, we blocked transcription by treating actinomycin D (5μg/ml) to the NIH 3T3 cells and harvested them every 2 hrs for each sample.

### Plasmids

To generate the pGL3 vector which contains promoters of DBP, we amplified the several forms of mouse Dbp promoters (UCSC Genome Browser on Mouse Dec. 2011 (GRCm38/mm10) Assembly, https://genome.ucsc.edu) with forward and reverse primers using Pfu polymerase (Solgent, Daejeon, South Korea). The PCR products were digested with Nhe1 and Hind3 and then inserted into the pGL3 basic vector, yielding pGL3 with the short form of Dbp promoter and pGL3 with the long form of Dbp promoter, which was confirmed by sequencing. Serial deletion constructs (P1, P2, P3, P4, P5, P6) and mutant constructs were cloned in the same way. Dbp-luciferase construct was kindly gifted by Ripperger (Stratmann et al., 2010). HA-hnRNP K and HA-hnRNP K (S284A/S353A) were kindly provided by Ronai (Habelhah et al., 2001).

### RNA extraction & quantitative PCR

Total RNA was extracted from NIH 3T3 and U2OS cells using the TRI Reagent (Molecular Research Center, Bio Science Technology). Next, we treated RQ1 RNase-free DNase (Promega) after RNA quantification to remove contaminated DNA according to the manufacturer’s instructions. RNA (1μg) was reverse-transcribed using ImProm-II™ (Promega) according to the manufacturer’s instructions. Endogenous mRNA levels were detected by quantitative real-time PCR using the StepOnePlus real-time PCR system (Applied Biosystems) with FastStart Universal SYBR Green Master (Roche), as described previously (65). Specific primer pairs for mouse Dbp (NM_016974.3), mouse pre-Dbp which primers targeted for intron regions (4), mouse Per1 (NM_011065.5 mouse Per2 (NM_011066.3), mouse Rev-erbα (NM_145434.4), mouse Hlf (NM_172563.3), mouse Tef (NM_017376.3), mouse Nfil3 (NM_017373.3) (18), mouse Bmal1 (NM_007489.4), mouse Per3 (NM_001289877.1) (20), mouse Cry1 (NM_007771.3), mouse Cry2 (NM_009963.4), human hnRNP K (NM_002140.4), and human DBP (NM_001352.4) were used for real-time PCR (the primer sequences are shown in Supplementary Table S1).

### Luciferase assay

Firefly and Renila luciferase activities were determined by using the Dual-Luciferase^®^ Reporter Assay System created by Promega (Promega Corporation, Wisconsin, USA) according to the manufacturer’s instructions. Normalized FLUC activity was determined as the ratio of Firefly to Renila activity.

### Real-time luminescence

5 × 10^5^ NIH 3T3 fibroblasts (ATCC) were plated in 35 mm dishes. Cells were transfected with 1 μg of each mDBP-luciferase construct using Lipofectamine 2000 (Invitrogen) according to the manufacturer’s instructions. After overnight culture, 100 nM dexamethasone was added to the cells and cells were incubated for 2 hr. The medium was then changed to DMEM without phenol red and 1 mM luciferin (Promega). A Lumicycle device (Actimetrics) kept in the 37°C incubator for recording measured luminescence for 3 or 4 days. Lumicycle analysis software (Actimetrics) was used for lumicycle data analysis.

### Generation of poly(C) motif region-deleted cell lines

We used the pSpCas9(BB)-2A-GFP (PX458) plasmid obtained from Addgene (Addgene, #48138) for Cas9-CRISPR experiments. We designed sgRNA sequences to delete poly(C) motif containing regions within the DBP promoter (66) by using the online CRISPR Design tool (http://crispr.mit.edu/). Sequences were as follows: Target 1 (Forward: 5’-CACCGCATTGGCCCGAAGTGGGTCA-3, Reverse: 5’-CTGACCCACTTCGGGCCAATGCAAA-3’) and Target 2 (Forward: 5’-CACCGGCGGGACACTGACCTATATT-3’, Reverse: 5’-CAATATAGGTCAGTGTCCCGCCAAA-3’). Then, genomic DNA sequencing was performed with the following primers: Forward, CTA GCT AGC CGA TAG CAC GCG CAA AGC CA; reverse, 5’-CCC AAG CTT GGC AAG AAC CAA TCA CGT CT-’ to confirm genomic deletion.

### ChIP assay

After fixation of 10^7^ cells of NIH 3T3 cells by 1% formaldehyde at RT for 10 min, 0.125 M glycine was treated for 10 min to quench the formaldehyde. Next, the fixed cells were suspended in 10 mM Tris-Cl, pH 8.0, 1 mM EDTA, 0.1% SDS, 0.5 mM PMSF, and protease inhibitor cocktail (Roche). Crosslinked cells were sonicated by a sonicator (VibraCell, Sonics, USA) with 20 cycles of sonication (30 sec) and resting (30 sec) at an amplitude of 8. Sonicated chromatin was analyzed by agarose gel electrophoresis to ensure that DNA fragment sizes did not exceed 500 bp. Anti-hnRNP K antibody (ab70492, Abcam) and Pol2 (8WG16, Abcam) was added to 750 ul of chromatin and incubated at 4°C for 2 hrs with rotation. 30 μl of Protein G magnetic beads (88848, Thermo) was added to the chromatin-antibody mixture and were incubated at 4°C overnight with rotation. Magnetic beads were washed twice with 1 ml of ChIP wash buffer 1 (10 mM Tris-Cl, pH 7.4, 1 mM EDTA, 0.1% SDS, 0.1% sodium deoxycholate, 1% Triton X-100) for 10 min, twice with 1 ml of ChIP wash buffer 2 (10 mM Tris-Cl, pH 7.4, 1 mM EDTA, 0.1% SDS, 0.1% sodium deoxycholate, 1% Triton X-100, 0.3 M NaCl) for 10 min, twice with 1ml of ChIP wash buffer 3 (10mM Tris-Cl, pH 7.4, 1mM EDTA, 0.25M LiCl, 0.5% NP-40, 0.5% sodium deoxycholate) for 10 min, once with 1ml of ChIP wash buffer 4 (10 mM Tris-Cl, pH 7.4, 1 mM EDTA, 0.2% Triton X-100) for 10 min, and once with 1 ml of 1X TE buffer (10 mM Tris-Cl, pH 7.4, 1 mM EDTA) for 10 min. Washed beads were pelleted on a magnetic stand and the clear supernatant was carefully removed and discarded. Beads were resuspended in 200μl of ChIP elution buffer (50mM Tris-Cl, pH 7.5, 10 mM EDTA, 1% SDS) and 5μl of proteinase K (Roche, Cat # 03 115 828 001) was added to beads suspension. The beads and proteinase K were mixed well by pipetting and were incubated at 65°C for 5 hrs. 200μl of phenol:chloroform: isoamyl alcohol (25: 24: 1) (Affymetrix, Cat. # 75831 400 ML) was added to beads suspension and mixed vigorously by vortexing for 30 sec and was centrifuged at 13,000 rpm for 15 min under room temperature. After centrifugation, the upper aqueous phase was transferred to a fresh 1.5ml tube and 2μl of glycogen solution (Roche, Cat. # 10901393001), 20μl of 3M sodium acetate, and 500μl of 100% ethanol were added and the solution was incubated at −20°C overnight. DNA was pelleted by centrifuging at 13,000 rpm for 30 min at 4°C. The supernatant was carefully removed and discarded. DNA pellet was washed with 700μl of 70% ethanol and DNA was pelleted by centrifuging at 13,000 rpm for 15 min at 4°C. The supernatant was carefully removed and discarded while the DNA pellet was air-dried for 15 min at room temperature. DNA was resuspended in 50μl of 1X TE buffer (10mM Tris-Cl, pH 7.5, 1mM EDTA) and used for quantification by quantitative PCR (the primer sequences are shown in Supplemental Table S2).

### ATAC-qPCR

ATAC-seq was performed as described (67) on four of wild-type NIH3T3 and four of poly(c) deleted NIH3T3. Libraries were generated using the Ad1_noMX, Ad2.13, Ad2.15, Ad2.16 and Ad2.17 barcoded primers from Buenrostro et al. (68) and were amplified for 6-7 total cycles. Libraries were purified with SPRI beads to remove remaining primer dimers. Libraries were quantitated by using Qubit.

### Subcellular fractionation

Hypotonic buffer (10mM HEPES (pH7.9), 10mM KCl, 1.5mM MgCl2, DTT 1.0mM, 0.2mM PMSF) was added to cells to resuspend them after cell harvest. Then, Lysis buffer (10mM HEPES (pH7.9), 10mM KCl, 1.5mM MgCl2, 1.0mM DTT, 0.2mM PMSF, 2.5 % NP-40) was added and the solution was centrifuged at 3500 rpm, 4°C for 4 min. Cytoplasmic fraction was extracted with 5 trials of freezing and thawing of the supernatant of cell lysate followed by centrifugation at 15000 rpm, 4°C, for 20 min. Next, to extract nuclear fraction, extraction buffer (20mM HEPES (pH7.9), 450mM NaCl, 1.5mM MgCl2, 1.0mM DTT, 0.2mM PMSF, 0.2mM EDTA (pH 8.0) was added to pellet of cell lysate and were centrifuged at 15000 rpm, 4°C, for 20 min after 5 trials of freezing and thawing. Each fraction was quantified by Bradford assay and were boiled at 95°C for 5 min with sample buffer (40% Glycerol, 240mM Tris/HCl pH 6.8, 8% SDS, 0.04% bromophenol blue, 5% beta-mercaptoethanol).

### Transient transfection and RNA interference

The Neon® Transfection System (Invitrogen) was used for transient transfection as recommended by the manufacturer, and Metafectene (Bionex) was used according to the manufacturer’s instruction. We transfected cells with 2 μg of luciferase reporter plasmid and maintained the cells in 6-well plates. For hnRNP K knockdown or overexpression and subsequent luciferase assays, we transfected cells with 1 μl of 20 μM siRNA or 1 μg of hnRNP K-overexpressing vectors (HA-pcDNA vectors) 12 hr after transfection of cells with 1 μg luciferase reporter plasmid. We also used specific small interfering RNAs (siRNAs) for hnRNP K siRNA pools (sihnRNP K1 sense: 5’aaaggaggcaagaauauua(dTdT)3’, antisense: 5’uaauauucuugccuccuuu(dTdT)3’, sihnRNP K2 sense: 5’gcuguggaaugcuuaaauu(dTdT)3’, antisense: 5’aauuuaagcauuccacagc(dTdT)3’, sihnRNP K3 sense: 5’cugaugagaugguugaauu(dTdT)3’, antisense: 5’aauucaaccaucucaucag(dTdT)3’, sihnRNP K4 sense: 5’ucaugaaucuggagcauca(dTdT)3’, antisense: 5’ugaugcuccagauucauga(dTdT)3’) (BIONEER, South Korea) to conduct knockdown of hnRNP K.

### Oligonucleotide pull-down assay and mass spectrometry analysis

We used biotinylated oligonucleotides (Solgent, South Korea) for oligonucleotide pull-down assays. The biotinylated oligonucleotides had the following sequences: biotinylated wild-type, 5’-CCG CCT CCA GCG CCT CCT CC-3’ and biotinylated mutant, 5’-AGG AAT ATA GCG ATT CAT AA-3’. After harvesting HEK293A cells, we lysed the cells using RIPA buffer (10 mM Tris-Cl, pH 7.4, 1 mM EDTA, 0.1% SDS, 0.1% sodium deoxycholate, 1% Triton X-100) and centrifuged them at 15,000 rpm for 15 min at 4°C. The supernatant was used as the cell lysate for the binding assay. First, we washed streptavidin agarose beaded resin (Thermo Scientific, Cat # 20349) and biotinylated oligonucleotides with RIPA buffer. Then we allowed binding of the streptavidin-agarose beaded resin to the biotinylated oligonucleotides and the cell lysate, GST, or GST-hnRNP K simultaneously for 1 hr to increase the specificity of pulldown. Next, we mixed streptavidin agarose beaded resin bound by biotinylated oligonucleotide with either precleared cell lysate, GST, or GST-hnRNP K to allow binding at 4°C for 4 hrs. The resin was then washed three times with RIPA buffer and boiled in sample buffer (40% glycerol, 240 mM Tris/HCl pH 6.8, 8% SDS, 0.04% bromophenol blue, 5% beta-mercaptoethanol) at 95°C for 10 min. Finally, SDS-PAGE was conducted. After SDS-PAGE, the gel was stained with Brilliant Blue R-250 (Sigma-Aldrich) for 30 min and destained with a solution of 30% methanol, 7% acetic acid, and 63% water. The gel was scanned for image acquisition and the band of interest was sent to Genomine, Inc. (Pohang, South Korea) for mass spectrometry. In-gel proteolytic digestion and protein fragment extraction were conducted and results were compared with the NCBI protein database using Mascot software to identify the protein.

### Protein extraction

GST and GST-hnRNP K were expressed in Escherichia coli BL21 (DE3) pLysS cells (Novagen) grown to an absorbance at 600nm (A600) of 0.6. Induction of GST, GST-hnRNP K was performed by 0.5mM isopropyl β-D-1-thiogalactopyranoside (IPTG) overnight at 18°C. Next, cells were resuspended in 20mM Tris–HCl (pH 7.5), 150mM NaCl, 1% Triton-X, 1mM dithiothreitol, and protease inhibitor cocktail (Roche) and were lysed by sonication. And, GST and GST-hnRNP K were purified using glutathione-sepharose 4B agarose beads (GE Healthcare Bio-Sciences) and were eluted in 50mM Tris-HCl (pH 8.0) and 20mM L-Glutathione reduced (GE Healthcare).

### Immunoblot assay

Immunoblotting was conducted by using polyclonal anti-hnRNP K (Abcam, ab70492), monoclonal anti-GST (Santa Cruz, sc-138), Polyclonal anti-FLAG (Cell signaling, #2368), polyclonal anti-14-3-3 (Santa Cruz, sc-1019) and, monoclonal anti-GAPDH (Santa Cruz, sc-47724) as primary antibodies. A SUPEX ECL solution kit (Neuronex) and LAS-4000 chemiluminescence detection system (FUJIFILM) were used after horseradish peroxidase-conjugated species-specific secondary antibodies (goat, Santa Cruz Biotechnology; guinea pig, Santa Cruz Biotechnology; mouse, Thermo Scientific; rabbit, Jackson ImmunoResearch Laboratories) reactions. And, Image Gauge (Fuji Film) was used for analysis of the acquired images according to the manufacturer’s instructions.

### MTT assay

NIH3T3 and U2OS cells at a density of 2×10^4^ cells per well in 96-well plates were transfected with siCon and sihnRNP K. After 48 hr, MTT (5 mg/ml) was added to each well followed by a 2-hr incubation at room temperature. Next, the media was removed and 100 μl DMSO was added to each well. Color development was allowed to proceed for 3 hr at room temperature in the dark. Finally, we measured the absorbance of each well at 570 nm using a NanoQuant spectrophotometer (Tecan).

### *Drosophila* stocks

Flies were raised on standard cornmeal-yeast-agar food at 25°C. *w*^1118^ (control), R18H11-Gal4, and UAS-hnRNPK^RNAi^ (T42540) lines were obtained from the Bloomington *Drosophila* Stock Center (BDSC). Two UAS-hnRNPK^RNAi^ lines (v2912, v106271) were obtained from the Vienna *Drosophila* RNAi Center. *Pdf*-Gal4 and *tim*-Gal4 lines were described previously. (61, 62)

### Behavioral analysis

Individual male flies were loaded into glass tubes containing 5% sucrose and 2% agar, entrained by five 12-hour light: 12-hour dark (LD) cycles, and then transferred to constant dark (DD) for 7 days at 25°C. Locomotor activities were continuously recorded in 1-min bins using the infrared beam-based *Drosophila* Activity Monitor (DAM) system (Trikinetics). Behavioral data were analyzed using the ClockLab software in MATLAB (Actimetrics). Circadian periods and the power of rhythmicity in DD locomotor rhythms were calculated in individual flies using the chi-square periodogram (the significance level was set to an alpha value of 0.05) and averaged per each genotype. Flies with the power of rhythmicity value less than 10 were defined as arrhythmic. Locomotor activity profiles were generated using a Microsoft Excel. Morning and evening indices anticipatory to lights-on and lights-off, respectively, were calculated as described previously (69). Dead flies were manually scored at the end of behavioral tests and their locomotor data were excluded from further analyses.

### Statistical analysis

Graphs are plotted as a mean±S.E.M which was illustrated by bars. Two-way ANOVA was used to conduct grouped analyses with multiple groups by Sidak’s multiple comparisons test. One-way ANOVA was used to conduct column analyses with more than two groups with Tukey’s multiple comparisons test analyzed with PRISM software.

## DATA AVAILABILITY

hnRNP K bound genes were available in Huarte et al. 2010 (Cell, 142, 409-419). MEME (Multiple Em for Motif Elicitation), MEME-ChIP, GOMo (Gene Ontology for Motifs) algorithms were available at http://meme-suite.org/

## ACKNOWLEDGEMENT

The authors thank Ze’ev Ronai (Sanford Burnham Prebys Medical Discovery Institute) for providing HA-hnRNP K WT, Jürgen A. Ripperger (University of Fribourg) for providing DBP-luciferase plasmid, and Sung-Hoon Kim (McGill University) for providing the GST-hnRNP K plasmid. This work was supported by the Bio & Medical Technology Development Program of the National Research Foundation (NRF) funded by the Korean government (MSIT) (No. 2018011982); “Cooperative Research Program for Agriculture Science & Technology Development (Project No. PJ01324801)” by Rural Development Administration; and BK21 Plus funded by the Ministry of Education, Republic of Korea (10Z20130012243).

## Author contributions

P. K. Kwon, K-H. Lee, J. Kim, S. Tae, H-M Kim, S. Ham, J-H Choi, Y-H Jeong, and S-W Kim conducted and analyzed the experiments; H. Yi and H-O Ku analyzed the lumicycle data; T-Y Roh, C. Lim, and K-T Kim designed the experiments; P. K. Kwon and K-T Kim wrote the paper.

