## Supplemental figures and tables for "Transcriptional activation of DBP by hnRNP K facilitates circadian rhythm"

### Slide 1
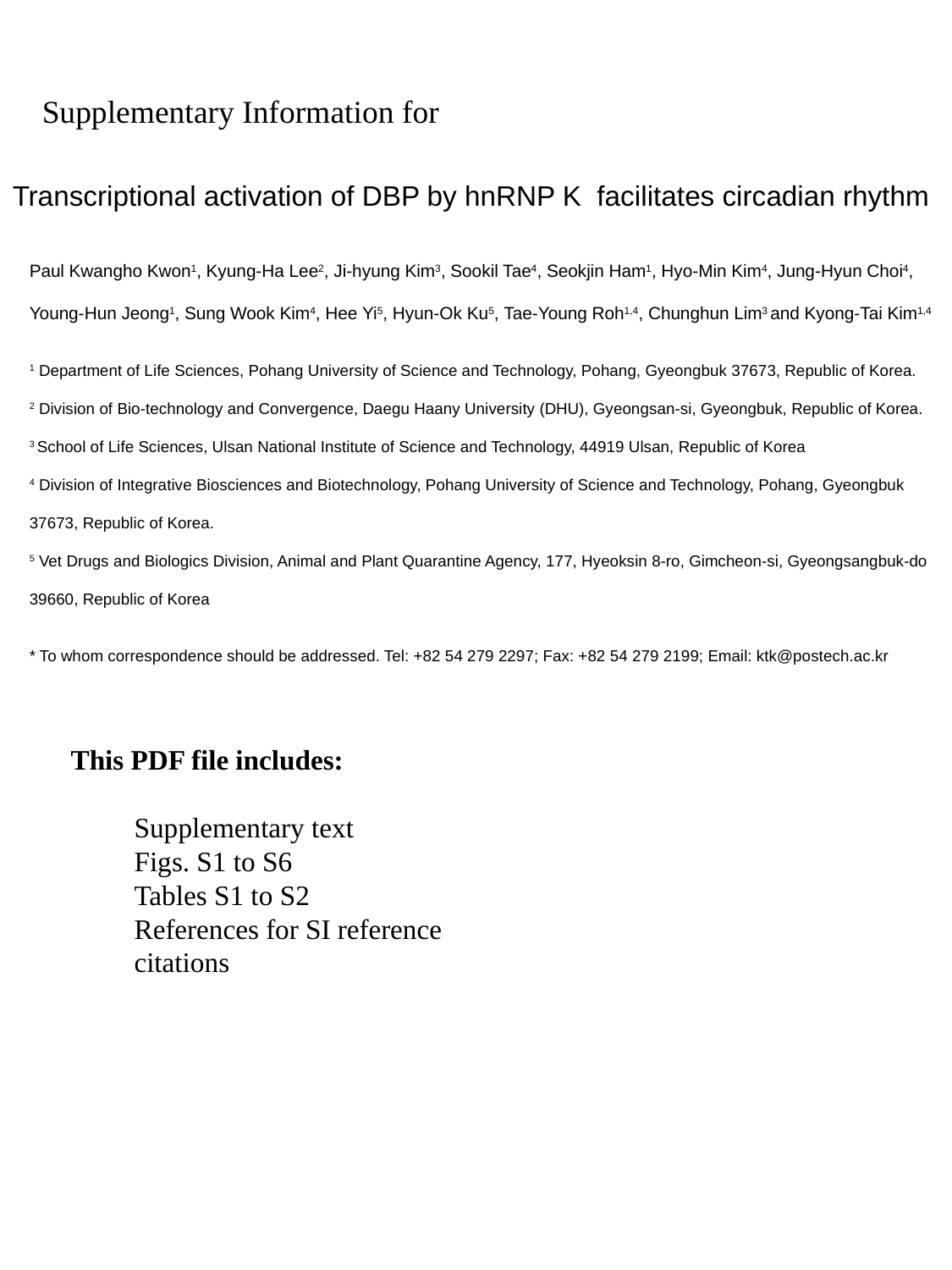

Supplementary Information for
Transcriptional activation of DBP by hnRNP K facilitates circadian rhythm
Paul Kwangho Kwon^1^, Kyung-Ha Lee^2^, Ji-hyung Kim^3^, Sookil Tae^4^, Seokjin Ham^1^, Hyo-Min Kim^4^, Jung‐Hyun Choi^4^, Young-Hun Jeong^1^, Sung Wook Kim^4^, Hee Yi^5^, Hyun-Ok Ku^5^, Tae-Young Roh^1,4^, Chunghun Lim^3^ and Kyong-Tai Kim^1,4^
^1^ Department of Life Sciences, Pohang University of Science and Technology, Pohang, Gyeongbuk 37673, Republic of Korea.
^2^ Division of Bio-technology and Convergence, Daegu Haany University (DHU), Gyeongsan-si, Gyeongbuk, Republic of Korea.
^3^ School of Life Sciences, Ulsan National Institute of Science and Technology, 44919 Ulsan, Republic of Korea
^4^ Division of Integrative Biosciences and Biotechnology, Pohang University of Science and Technology, Pohang, Gyeongbuk 37673, Republic of Korea.
^5^ Vet Drugs and Biologics Division, Animal and Plant Quarantine Agency, 177, Hyeoksin 8-ro, Gimcheon-si, Gyeongsangbuk-do 39660, Republic of Korea
This PDF file includes:
Supplementary text
Figs. S1 to S6
Tables S1 to S2
References for SI reference citations

### Slide 2
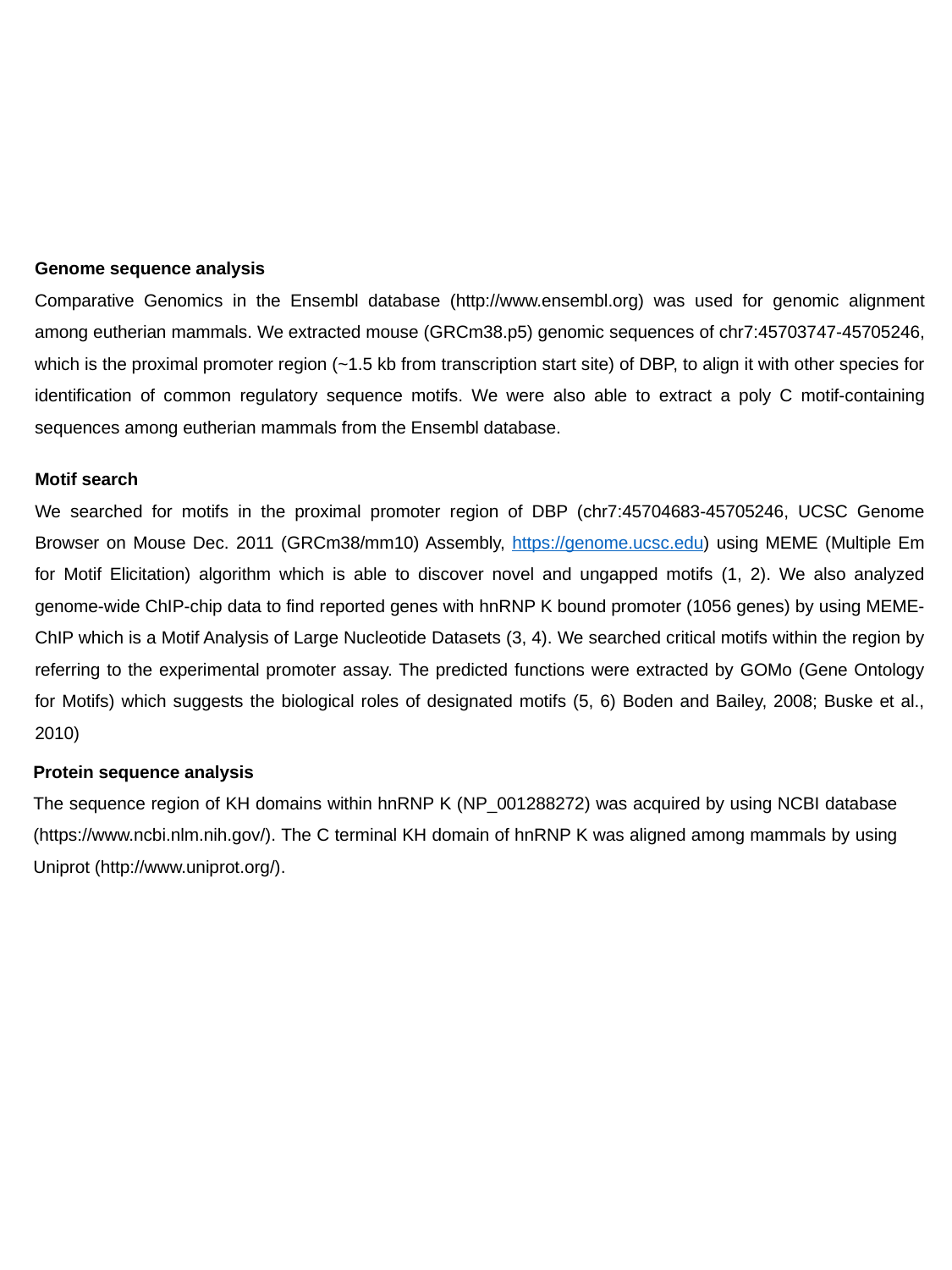

#
Genome sequence analysis
Comparative Genomics in the Ensembl database (http://www.ensembl.org) was used for genomic alignment among eutherian mammals. We extracted mouse (GRCm38.p5) genomic sequences of chr7:45703747-45705246, which is the proximal promoter region (~1.5 kb from transcription start site) of DBP, to align it with other species for identification of common regulatory sequence motifs. We were also able to extract a poly C motif-containing sequences among eutherian mammals from the Ensembl database.
Motif search
We searched for motifs in the proximal promoter region of DBP (chr7:45704683-45705246, UCSC Genome Browser on Mouse Dec. 2011 (GRCm38/mm10) Assembly, https://genome.ucsc.edu) using MEME (Multiple Em for Motif Elicitation) algorithm which is able to discover novel and ungapped motifs (1, 2). We also analyzed genome-wide ChIP-chip data to find reported genes with hnRNP K bound promoter (1056 genes) by using MEME-ChIP which is a Motif Analysis of Large Nucleotide Datasets (3, 4). We searched critical motifs within the region by referring to the experimental promoter assay. The predicted functions were extracted by GOMo (Gene Ontology for Motifs) which suggests the biological roles of designated motifs (5, 6) Boden and Bailey, 2008; Buske et al., 2010)
Protein sequence analysis
The sequence region of KH domains within hnRNP K (NP_001288272) was acquired by using NCBI database (https://www.ncbi.nlm.nih.gov/). The C terminal KH domain of hnRNP K was aligned among mammals by using Uniprot (http://www.uniprot.org/).

### Slide 3
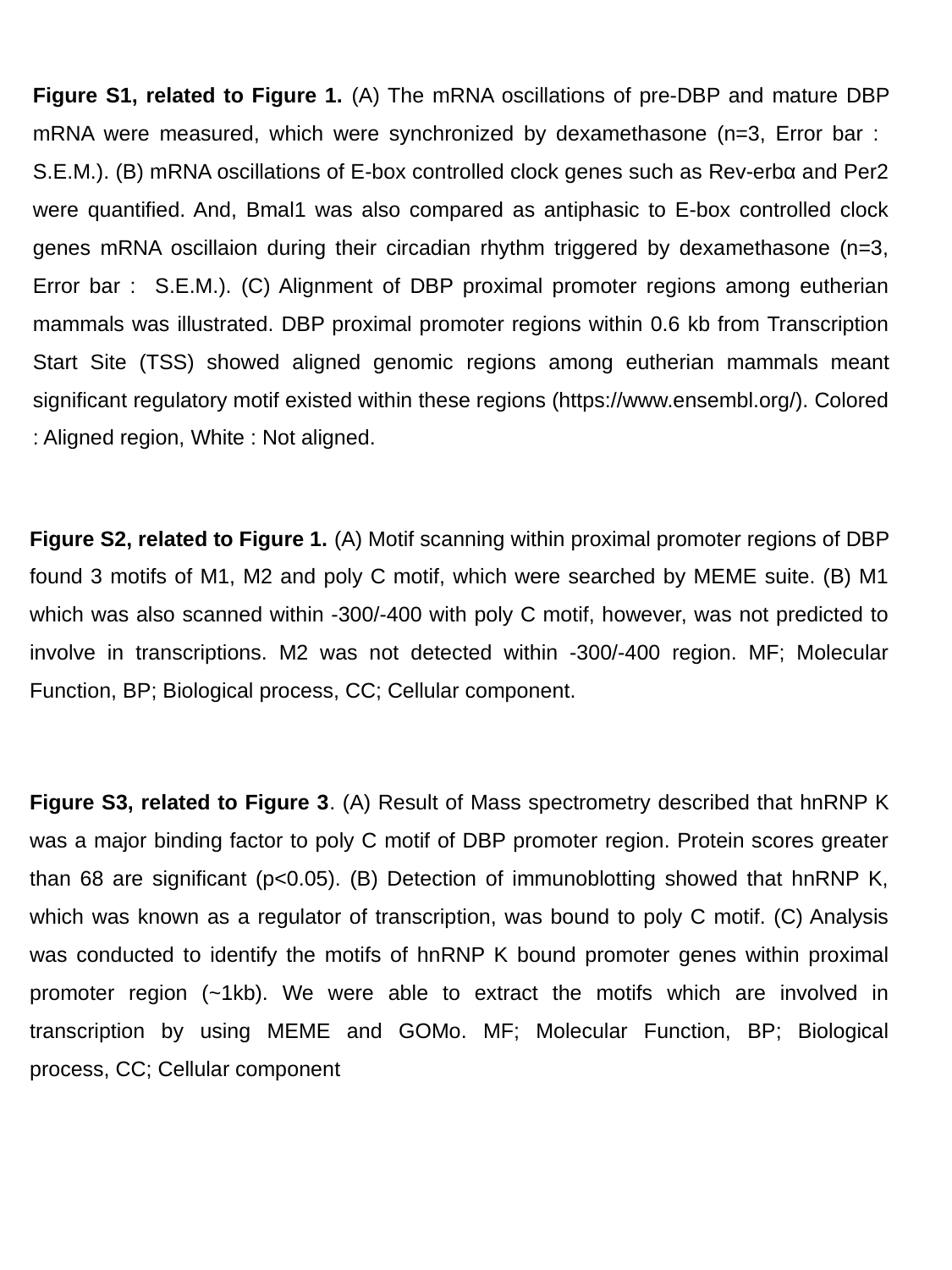

Figure S1, related to Figure 1. (A) The mRNA oscillations of pre-DBP and mature DBP mRNA were measured, which were synchronized by dexamethasone (n=3, Error bar : S.E.M.). (B) mRNA oscillations of E-box controlled clock genes such as Rev-erbα and Per2 were quantified. And, Bmal1 was also compared as antiphasic to E-box controlled clock genes mRNA oscillaion during their circadian rhythm triggered by dexamethasone (n=3, Error bar : S.E.M.). (C) Alignment of DBP proximal promoter regions among eutherian mammals was illustrated. DBP proximal promoter regions within 0.6 kb from Transcription Start Site (TSS) showed aligned genomic regions among eutherian mammals meant significant regulatory motif existed within these regions (https://www.ensembl.org/). Colored : Aligned region, White : Not aligned.
Figure S2, related to Figure 1. (A) Motif scanning within proximal promoter regions of DBP found 3 motifs of M1, M2 and poly C motif, which were searched by MEME suite. (B) M1 which was also scanned within -300/-400 with poly C motif, however, was not predicted to involve in transcriptions. M2 was not detected within -300/-400 region. MF; Molecular Function, BP; Biological process, CC; Cellular component.
Figure S3, related to Figure 3. (A) Result of Mass spectrometry described that hnRNP K was a major binding factor to poly C motif of DBP promoter region. Protein scores greater than 68 are significant (p<0.05). (B) Detection of immunoblotting showed that hnRNP K, which was known as a regulator of transcription, was bound to poly C motif. (C) Analysis was conducted to identify the motifs of hnRNP K bound promoter genes within proximal promoter region (~1kb). We were able to extract the motifs which are involved in transcription by using MEME and GOMo. MF; Molecular Function, BP; Biological process, CC; Cellular component

### Slide 4
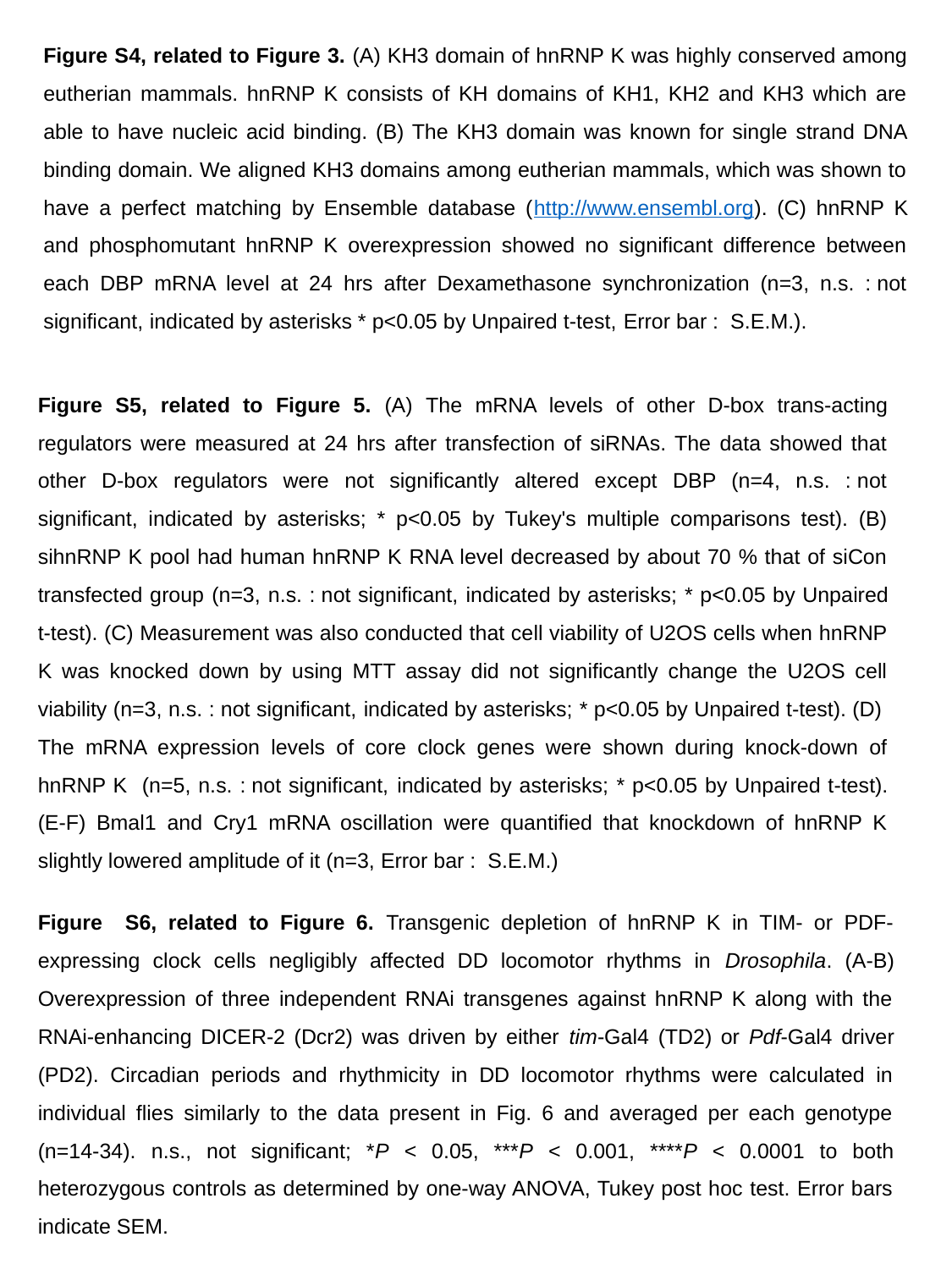

Figure S4, related to Figure 3. (A) KH3 domain of hnRNP K was highly conserved among eutherian mammals. hnRNP K consists of KH domains of KH1, KH2 and KH3 which are able to have nucleic acid binding. (B) The KH3 domain was known for single strand DNA binding domain. We aligned KH3 domains among eutherian mammals, which was shown to have a perfect matching by Ensemble database (http://www.ensembl.org). (C) hnRNP K and phosphomutant hnRNP K overexpression showed no significant difference between each DBP mRNA level at 24 hrs after Dexamethasone synchronization (n=3, n.s. : not significant, indicated by asterisks * p<0.05 by Unpaired t-test, Error bar : S.E.M.).
Figure S5, related to Figure 5. (A) The mRNA levels of other D-box trans-acting regulators were measured at 24 hrs after transfection of siRNAs. The data showed that other D-box regulators were not significantly altered except DBP (n=4, n.s. : not significant, indicated by asterisks; * p<0.05 by Tukey's multiple comparisons test). (B) sihnRNP K pool had human hnRNP K RNA level decreased by about 70 % that of siCon transfected group (n=3, n.s. : not significant, indicated by asterisks; * p<0.05 by Unpaired t-test). (C) Measurement was also conducted that cell viability of U2OS cells when hnRNP K was knocked down by using MTT assay did not significantly change the U2OS cell viability (n=3, n.s. : not significant, indicated by asterisks; * p<0.05 by Unpaired t-test). (D) The mRNA expression levels of core clock genes were shown during knock-down of hnRNP K (n=5, n.s. : not significant, indicated by asterisks; * p<0.05 by Unpaired t-test). (E-F) Bmal1 and Cry1 mRNA oscillation were quantified that knockdown of hnRNP K slightly lowered amplitude of it (n=3, Error bar : S.E.M.)
Figure S6, related to Figure 6. Transgenic depletion of hnRNP K in TIM- or PDF-expressing clock cells negligibly affected DD locomotor rhythms in Drosophila. (A-B) Overexpression of three independent RNAi transgenes against hnRNP K along with the RNAi-enhancing DICER-2 (Dcr2) was driven by either tim-Gal4 (TD2) or Pdf-Gal4 driver (PD2). Circadian periods and rhythmicity in DD locomotor rhythms were calculated in individual flies similarly to the data present in Fig. 6 and averaged per each genotype (n=14-34). n.s., not significant; *P < 0.05, ***P < 0.001, ****P < 0.0001 to both heterozygous controls as determined by one-way ANOVA, Tukey post hoc test. Error bars indicate SEM.

### Slide 5
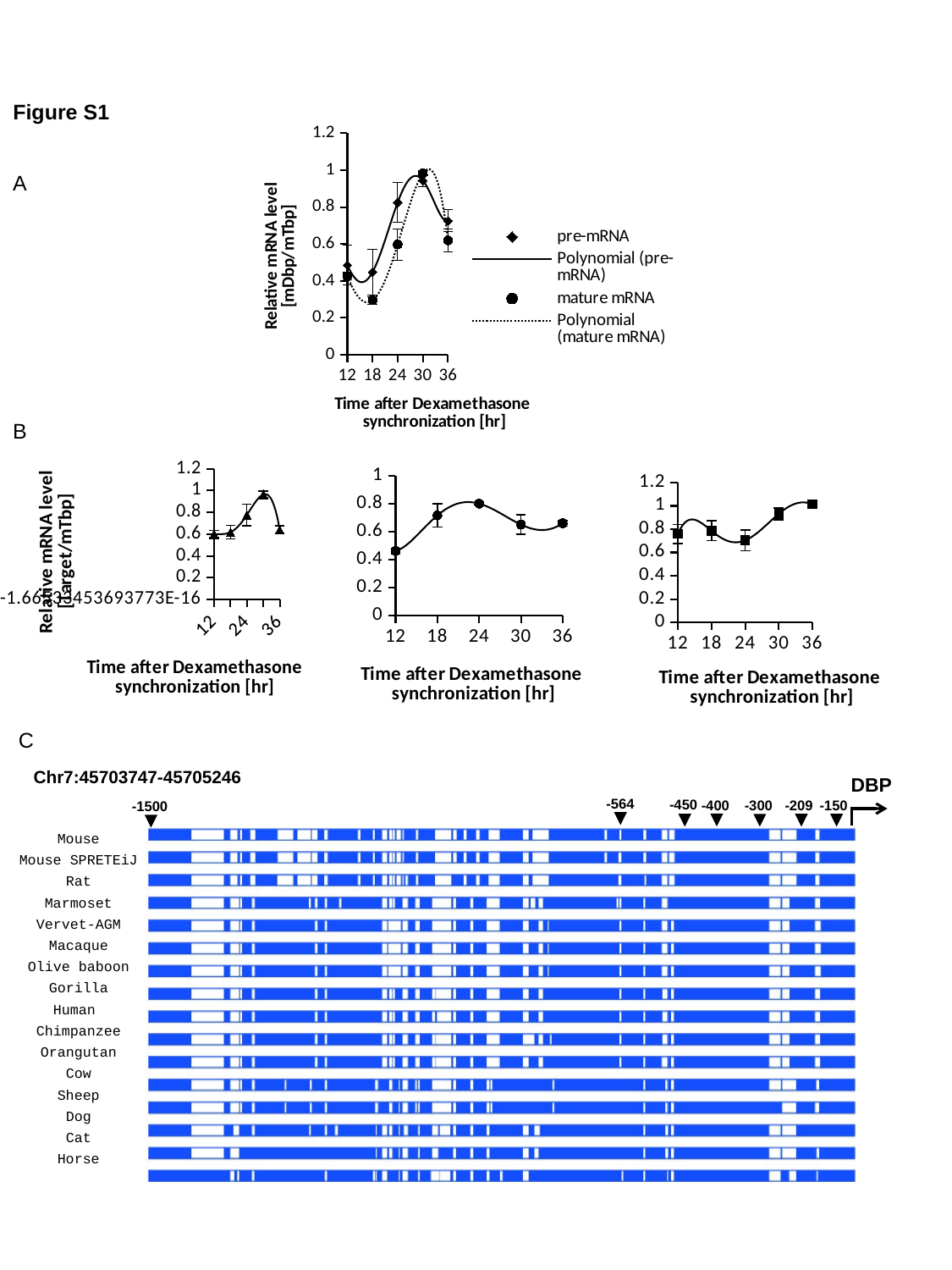

Figure S1
###### Chart
| Category | pre-mRNA | mature mRNA |
|---|---|---|A
B
###### Chart
| Category | Per2 |
|---|---|
###### Chart
| Category | Rev-erbα |
|---|---|
###### Chart
| Category | Bmal1 |
|---|---|C
 Chr7:45703747-45705246
DBP
-564
-450
-300
-209
-150
-1500
Mouse
Mouse SPRETEiJ
Rat
Marmoset
Vervet-AGM
Macaque
Olive baboon
Gorilla
Human
Chimpanzee
Orangutan
Cow
Sheep
Dog
Cat
Horse
-400

### Slide 6
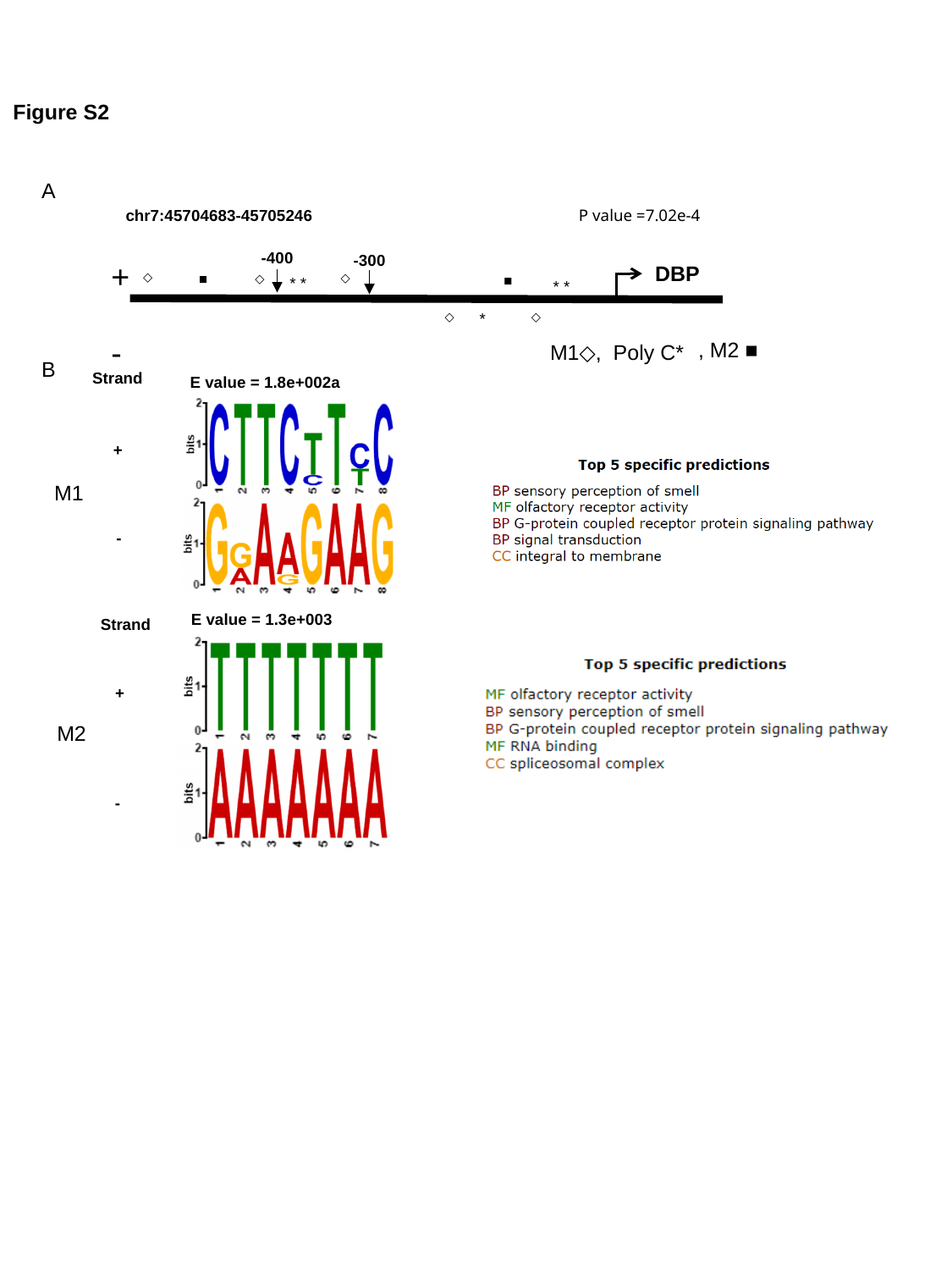

Figure S2
A
P value =7.02e-4
chr7:45704683-45705246
-400
-300
+
-
DBP
◇
■
◇
◇
■
* *
 * *
◇
*
◇
M1◇, Poly C*
, M2 ■
B
Strand
  E value = 1.8e+002a
+
M1
-
  E value = 1.3e+003
Strand
+
M2
-

### Slide 7
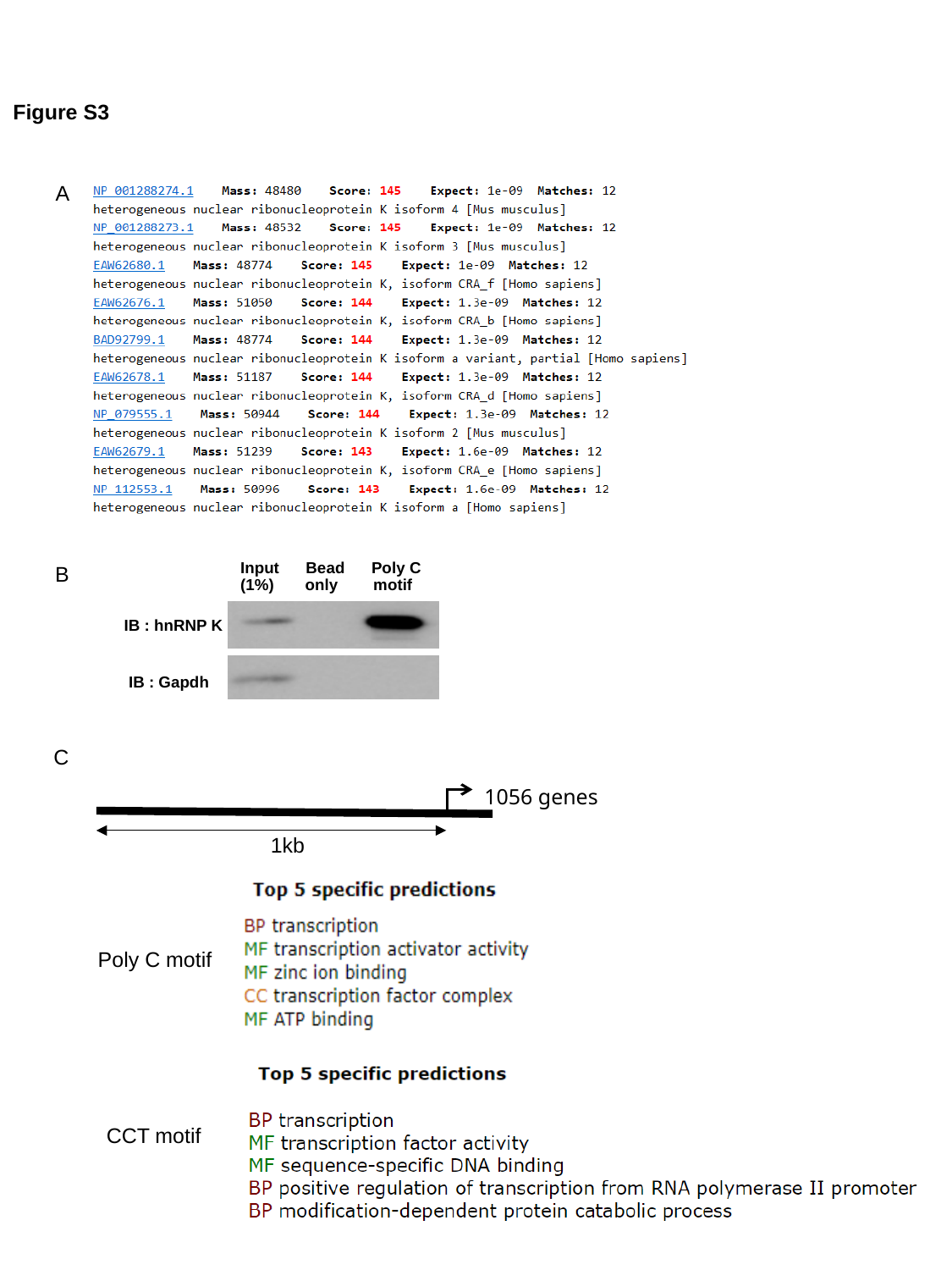

Figure S3
A
Input Bead Poly C
(1%) only motif
IB : hnRNP K
 IB : Gapdh
B
C
1056 genes
1kb
Poly C motif
CCT motif

### Slide 8
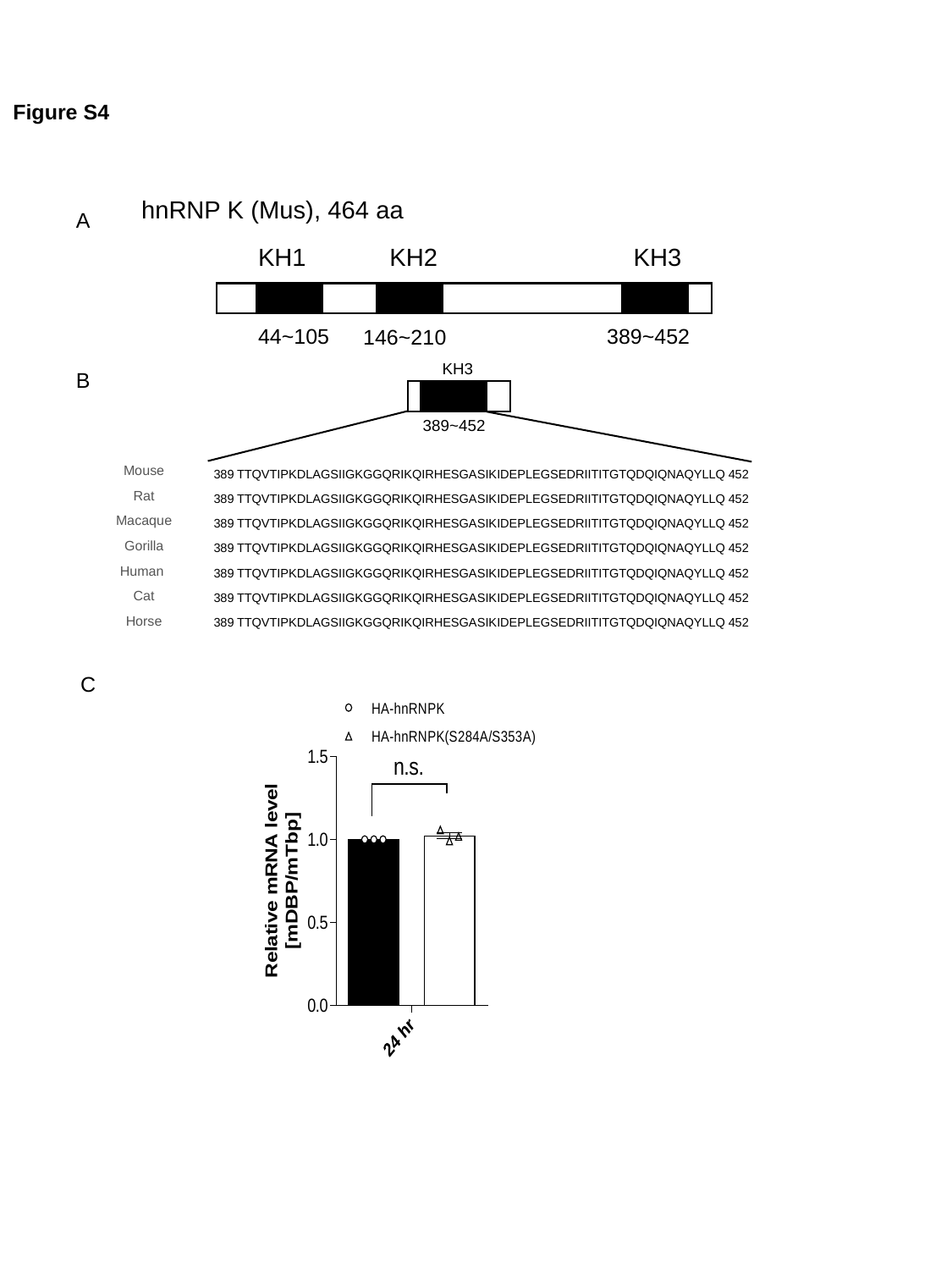

Figure S4
hnRNP K (Mus), 464 aa
 KH1 KH2	 KH3
A
44~105
389~452
146~210
	 	 KH3
B
389~452
389 TTQVTIPKDLAGSIIGKGGQRIKQIRHESGASIKIDEPLEGSEDRIITITGTQDQIQNAQYLLQ 452
389 TTQVTIPKDLAGSIIGKGGQRIKQIRHESGASIKIDEPLEGSEDRIITITGTQDQIQNAQYLLQ 452
389 TTQVTIPKDLAGSIIGKGGQRIKQIRHESGASIKIDEPLEGSEDRIITITGTQDQIQNAQYLLQ 452
389 TTQVTIPKDLAGSIIGKGGQRIKQIRHESGASIKIDEPLEGSEDRIITITGTQDQIQNAQYLLQ 452
389 TTQVTIPKDLAGSIIGKGGQRIKQIRHESGASIKIDEPLEGSEDRIITITGTQDQIQNAQYLLQ 452
389 TTQVTIPKDLAGSIIGKGGQRIKQIRHESGASIKIDEPLEGSEDRIITITGTQDQIQNAQYLLQ 452
389 TTQVTIPKDLAGSIIGKGGQRIKQIRHESGASIKIDEPLEGSEDRIITITGTQDQIQNAQYLLQ 452
Mouse
Rat
Macaque
Gorilla
Human
Cat
Horse
C

### Slide 9
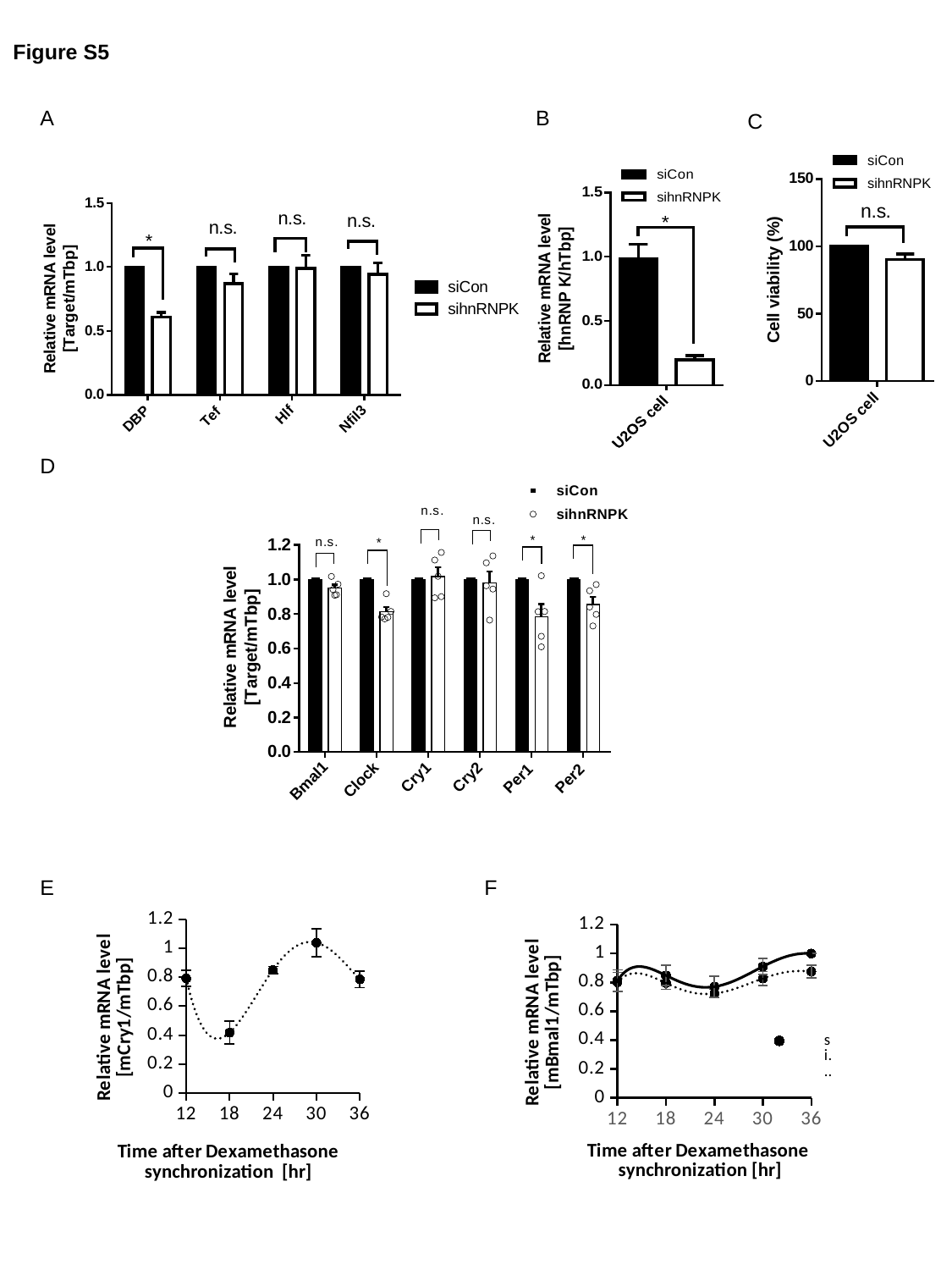

Figure S5
A
B
C
D
E
F
###### Chart
| Category | siCon | sihnRNP K |
|---|---|---|
###### Chart
| Category | siCon | sihnRNP K |
|---|---|---|

### Slide 10
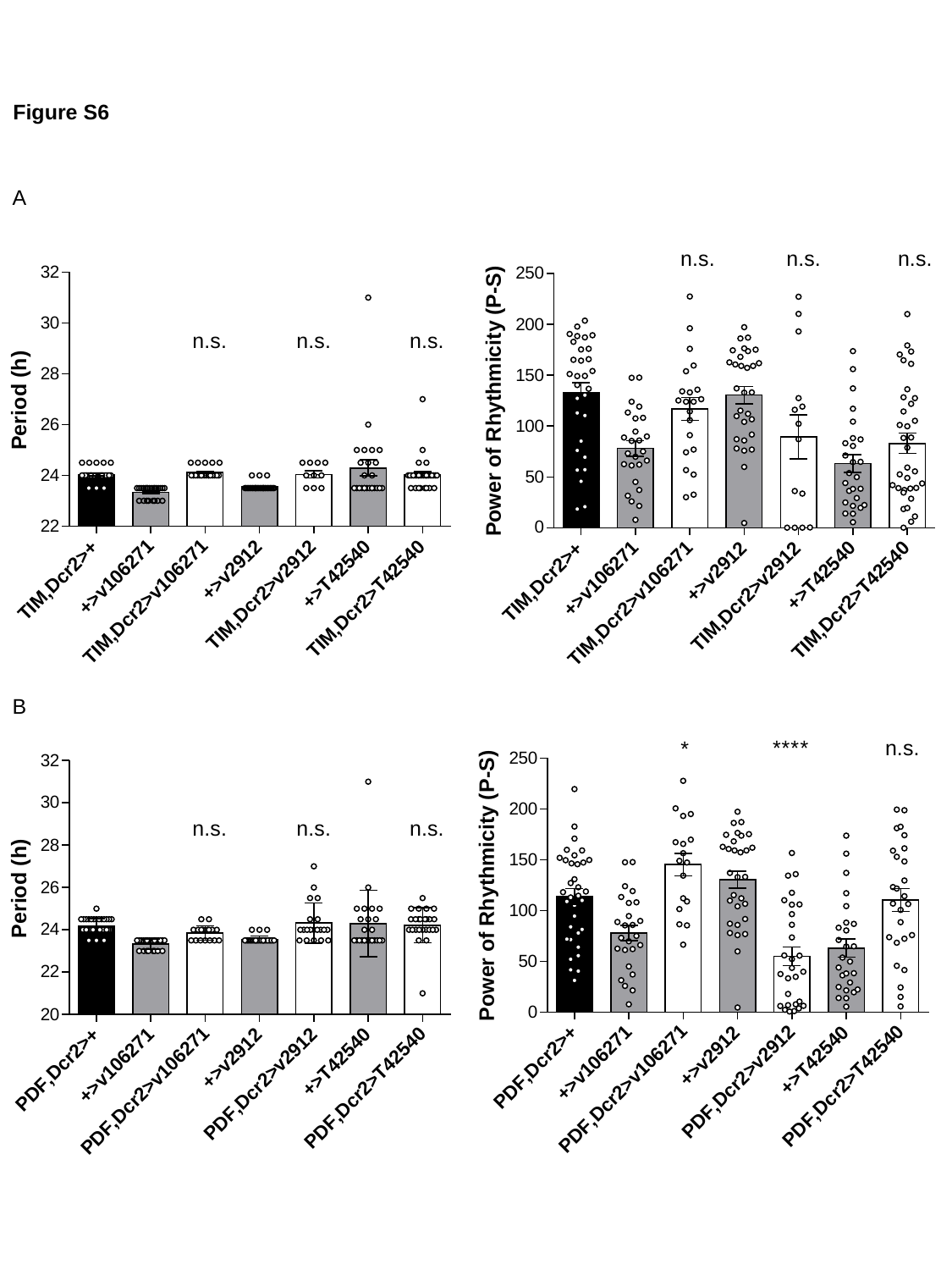

Figure S6
A
B

### Slide 11
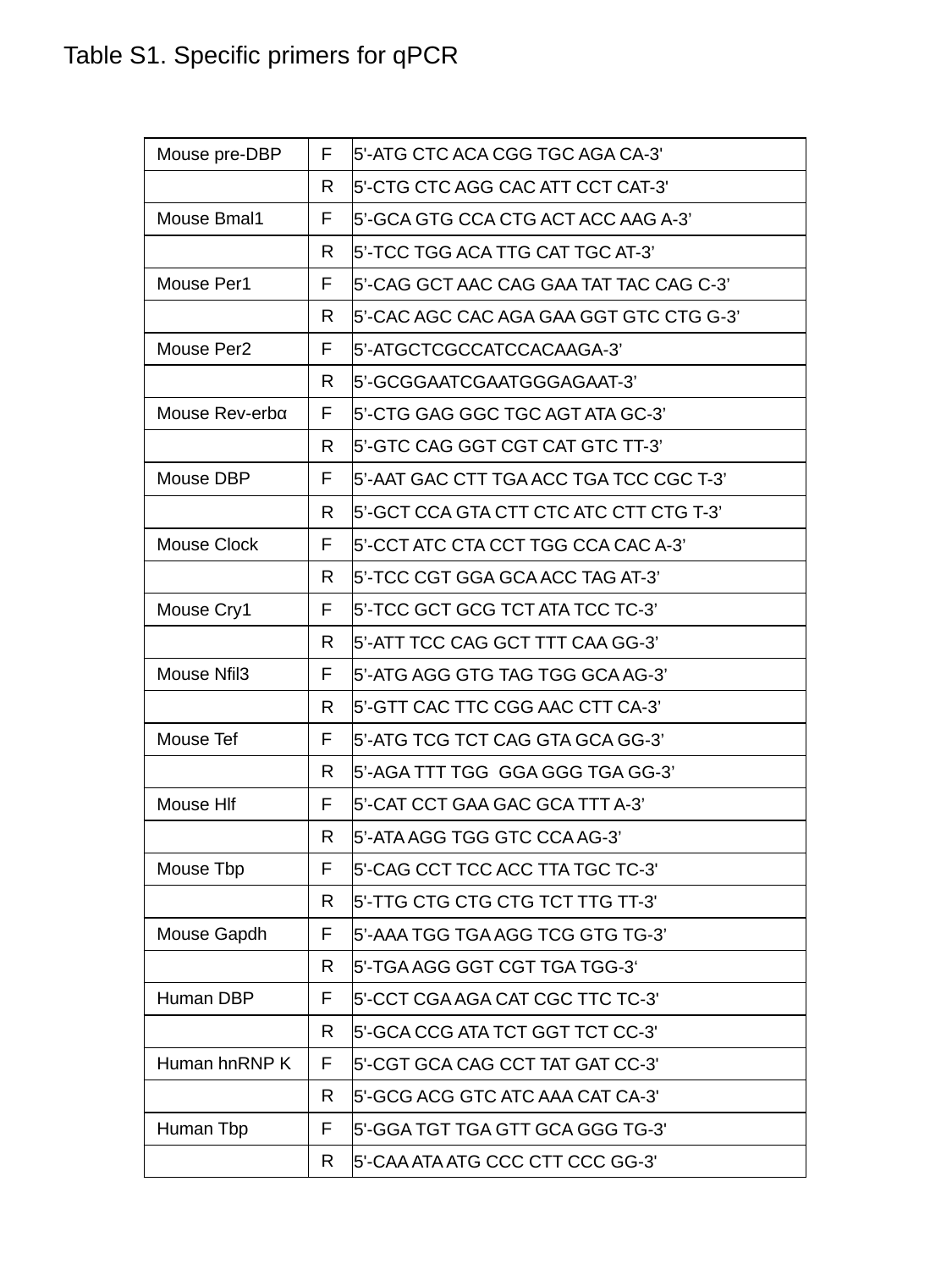

Table S1. Specific primers for qPCR
| Mouse pre-DBP | F | 5'-ATG CTC ACA CGG TGC AGA CA-3' |
| --- | --- | --- |
| | R | 5'-CTG CTC AGG CAC ATT CCT CAT-3' |
| Mouse Bmal1 | F | 5’-GCA GTG CCA CTG ACT ACC AAG A-3’ |
| | R | 5’-TCC TGG ACA TTG CAT TGC AT-3’ |
| Mouse Per1 | F | 5’-CAG GCT AAC CAG GAA TAT TAC CAG C-3’ |
| | R | 5’-CAC AGC CAC AGA GAA GGT GTC CTG G-3’ |
| Mouse Per2 | F | 5’-ATGCTCGCCATCCACAAGA-3’ |
| | R | 5’-GCGGAATCGAATGGGAGAAT-3’ |
| Mouse Rev-erbα | F | 5’-CTG GAG GGC TGC AGT ATA GC-3’ |
| | R | 5’-GTC CAG GGT CGT CAT GTC TT-3’ |
| Mouse DBP | F | 5’-aat gac ctt tga acc tga tcc cgc t-3’ |
| | R | 5’-gct cca gta ctt ctc atc ctt ctg t-3’ |
| Mouse Clock | F | 5’-CCT ATC CTA CCT TGG CCA CAC A-3’ |
| | R | 5’-TCC CGT GGA GCA ACC TAG AT-3’ |
| Mouse Cry1 | F | 5’-TCC GCT GCG TCT ATA TCC TC-3’ |
| | R | 5’-att tcc cag gct ttt caa gg-3’ |
| Mouse Nfil3 | F | 5’-atg agg gtg tag tgg gca ag-3’ |
| | R | 5’-gtt cac ttc cgg aac ctt ca-3’ |
| Mouse Tef | F | 5’-atg tcg tct cag gta gca gg-3’ |
| | R | 5’-aga ttt tgg gga ggg tga gg-3’ |
| Mouse Hlf | F | 5’-CAT CCT GAA GAC GCA TTT A-3’ |
| | R | 5’-ATA AGG TGG GTC CCA AG-3’ |
| Mouse Tbp | F | 5'-CAG CCT TCC ACC TTA TGC TC-3' |
| | R | 5'-TTG CTG CTG CTG TCT TTG TT-3' |
| Mouse Gapdh | F | 5’-AAA TGG TGA AGG TCG GTG TG-3’ |
| | R | 5'-TGA AGG GGT CGT TGA TGG-3‘ |
| Human DBP | F | 5'-CCT CGA AGA CAT CGC TTC TC-3' |
| | R | 5'-GCA CCG ATA TCT GGT TCT CC-3' |
| Human hnRNP K | F | 5'-CGT GCA CAG CCT TAT GAT CC-3' |
| | R | 5'-GCG ACG GTC ATC AAA CAT CA-3' |
| Human Tbp | F | 5'-GGA TGT TGA GTT GCA GGG TG-3' |
| | R | 5'-CAA ATA ATG CCC CTT CCC GG-3' |

### Slide 12
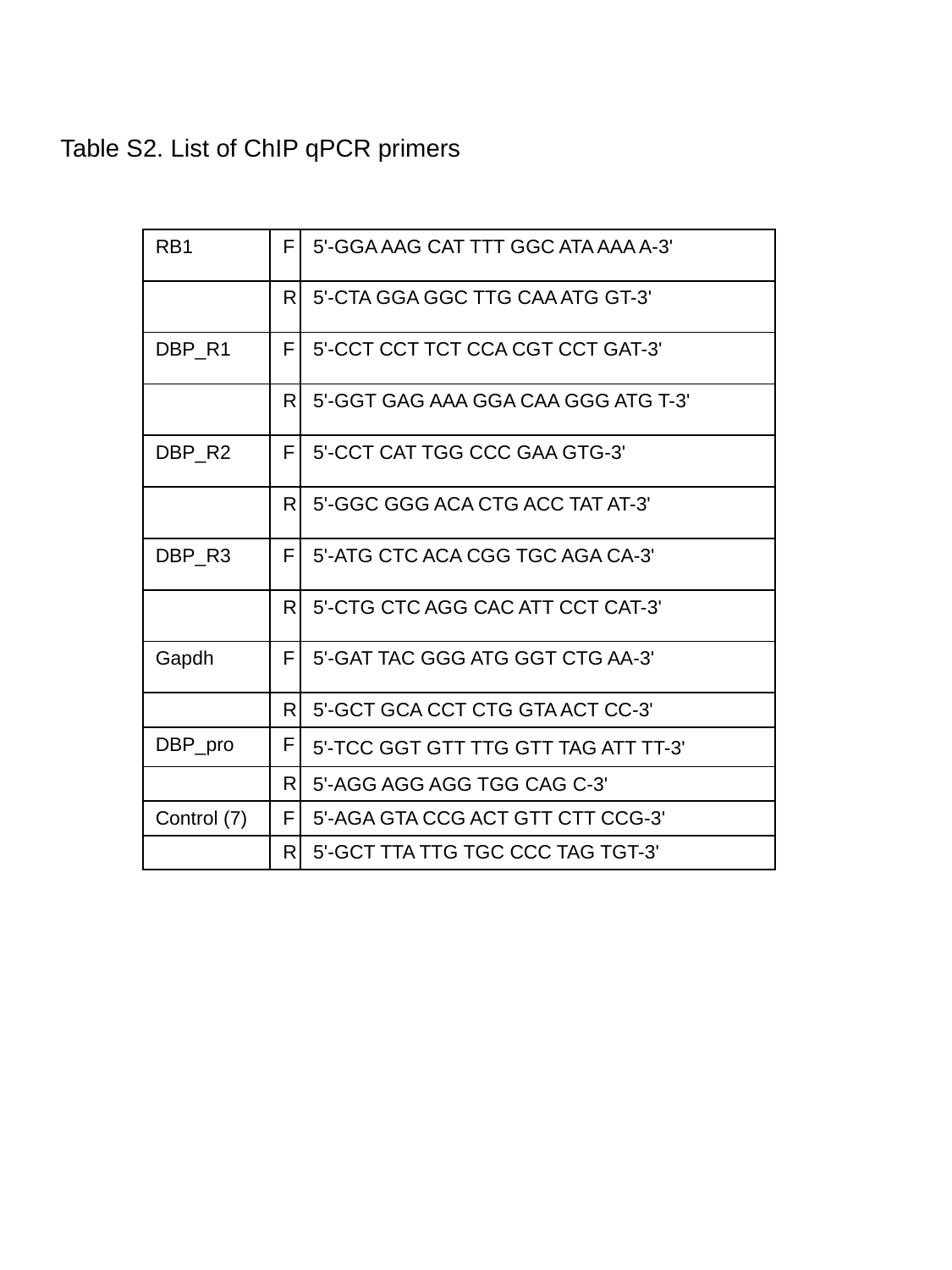

#### Table S2. List of ChIP qPCR primers
| RB1 | F | 5'-GGA AAG CAT TTT GGC ATA AAA A-3' |
| --- | --- | --- |
| | R | 5'-CTA GGA GGC TTG CAA ATG GT-3' |
| DBP\_R1 | F | 5'-CCT CCT TCT CCA CGT CCT GAT-3' |
| | R | 5'-GGT GAG AAA GGA CAA GGG ATG T-3' |
| DBP\_R2 | F | 5'-CCT CAT TGG CCC GAA GTG-3' |
| | R | 5'-GGC GGG ACA CTG ACC TAT AT-3' |
| DBP\_R3 | F | 5'-ATG CTC ACA CGG TGC AGA CA-3' |
| | R | 5'-CTG CTC AGG CAC ATT CCT CAT-3' |
| Gapdh | F | 5'-GAT TAC GGG ATG GGT CTG AA-3' |
| | R | 5'-GCT GCA CCT CTG GTA ACT CC-3' |
| DBP\_pro | F | 5'-TCC GGT GTT TTG GTT TAG ATT TT-3' |
| | R | 5'-AGG AGG AGG TGG CAG C-3' |
| Control (7) | F | 5'-AGA GTA CCG ACT GTT CTT CCG-3' |
| | R | 5'-GCT TTA TTG TGC CCC TAG TGT-3' |

### Slide 13
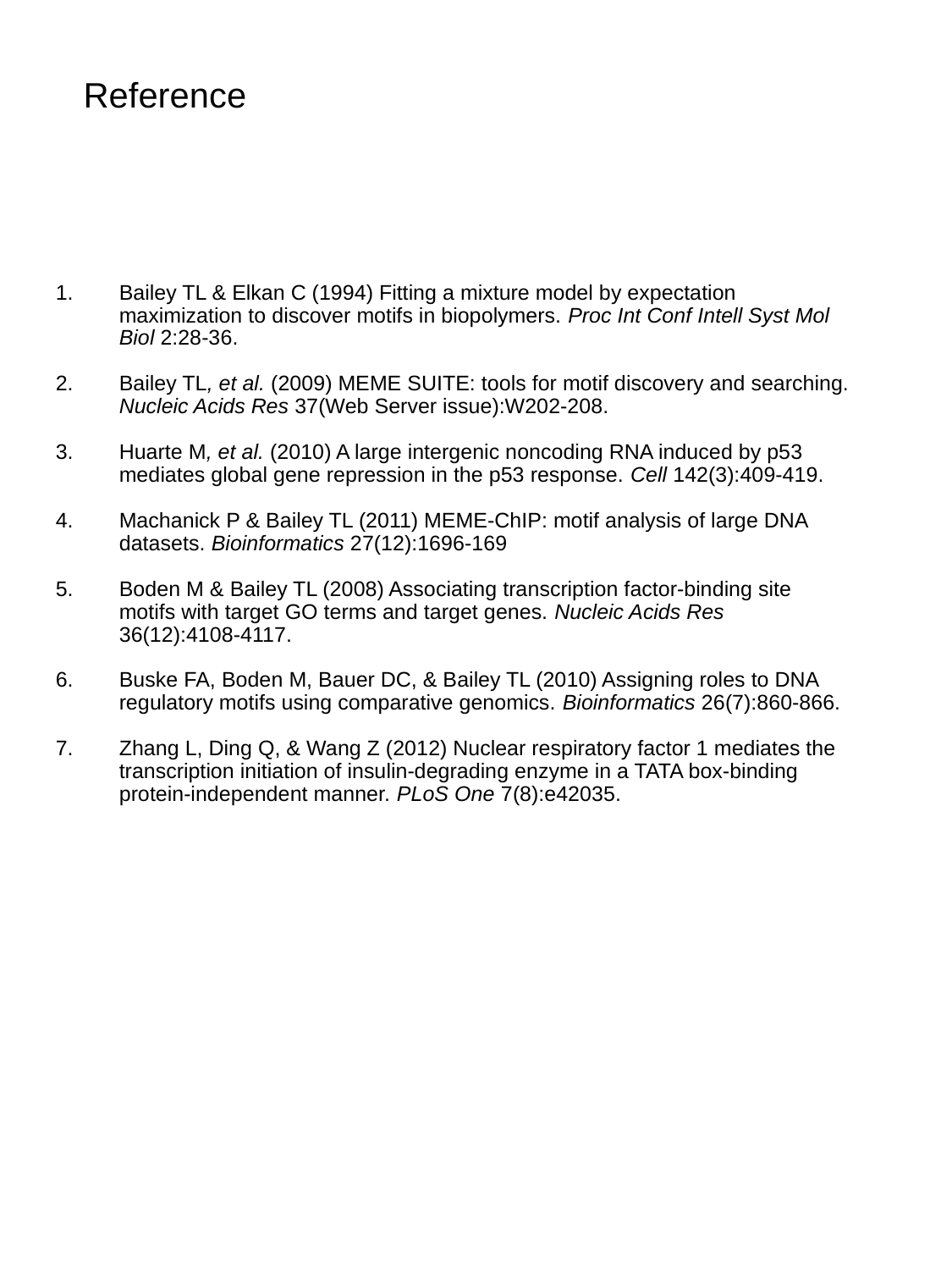
